# Sparse Autoencoders Reveal Interpretable Features in Single-Cell Foundation Models

**DOI:** 10.1101/2025.10.22.681631

**Authors:** Flavia Pedrocchi, Florian Barkmann, Amir Joudaki, Valentina Boeva

## Abstract

Single-cell foundation models (scFMs) hold promise for applications in cell type annotation, data integration, and prediction of the effects of cell perturbations, but their internal mechanisms remain poorly understood. We investigate the structure of these models by training sparse autoencoders (SAEs) on the hidden representations of three widely used scFMs: scGPT, scFoundation, and Geneformer.The learned features reveal diverse and complex biological and technical signals, which emerge even in pre-trained models. We also observe that the encoding of this information differs between scFMs with distinct training protocols and architectures. Finally, we demonstrate that SAE-derived features are functionally related to model behavior and can be intervened upon. Suppressing batch-associated features reduces unwanted technical variation and improves data integration while preserving the core biological signal. Activating drug-encoding features steers control cells toward drug-perturbed states in a concentration-dependent manner. These findings provide a path toward more interpretable and controllable single-cell foundation models.

## 1. Introduction

Single-cell foundation models (scFMs) have emerged as a powerful tool in computational biology to analyze cellular states and behavior (Bunne et al., 2024; Szałata et al., 2024). These models are typically trained on large-scale single-cell RNA sequencing datasets using self-supervised learning objectives (Ahlmann-Eltze et al., 2026). They can then be applied to several downstream tasks such as cell type annotation (Rosen et al., 2023; Heimberg et al., 2025), batch integration (He et al., 2025; Cui et al., 2024) or perturbation prediction, including drug response (Hao et al., 2024; Theus et al., 2024), and gene perturbation response (Adduri et al., 2025). The promise of scFMs lies in their ability to capture complex, non-linear relationships in high-dimensional transcriptomics data and generalize across different cell types, tissues, and experimental conditions.

Single-cell foundation models have found growing utility in single-cell analysis, yet many remain difficult to interpret. Like other large models, they often function as black boxes, making it unclear how their predictions are made (Fu et al., 2024). As these models are increasingly explored for generating biological insights, understanding what drives their predictions becomes important. This lack of transparency is particularly problematic given that their architectures are largely inherited from natural language processing models, with only minimal adaptation to the biological context (Theodoris et al., 2023; Yang et al., 2022). Moreover, despite their theoretical promise and computational complexity, several limitations of scFMs have been identified. Recent benchmarking studies have revealed mixed performance, with some finding that linear models can match or outperform scFMs in tasks such as cell type classification (Kedzierska et al., 2023) and perturbation response prediction (Ahlmann-Eltze et al., 2024; Csendes et al., 2024; Kernfeld et al., 2024), though performance appears to be task- and dataset-dependent. Additionally, scFMs often require fine-tuning to achieve practical usability (Liu et al., 2024; Steiner et al., 2025; Ovcharenko et al., 2025). Gaining a deeper understanding of how these models make predictions is essential for guiding future improvements in their design and training strategies.

Recent advances in mechanistic interpretability have introduced sparse autoencoders as a powerful technique for decomposing learned representations into interpretable, sparsely activated features that correspond to meaningful concepts (Cunningham et al., 2023). This approach has yielded insights into the internal workings of transformer-based language models (Templeton et al., 2024), DNA language models (Guan et al., 2025), and protein language models (Simon & Zou, 2024; Adams et al., 2025; Gujral et al., 2025). In this work, we utilize sparse autoencoders to understand the internal representations of scFMs, aiming to uncover the biological features these models learn, determine whether their representations align with biological knowledge, and assess the extent to which they encode technical artifacts.

### Our main contributions are

1. We show that pre-trained scFM can have a complex and meaningful understanding of cell biology.
2. We show how model architectures and training procedures can affect the encoding of information within the model.
3. We characterize how pre-trained scFMs represent technical variation alongside biological information, finding that cell type features often show study-specific patterns rather than consistent activation across all datasets.
4. We demonstrate that SAE-derived features are functionally related to model behavior and can be intervened upon to improve batch integration in scFMs.
5. We show that activating SAE-derived features can induce perturbation-associated cellular states learned by the scFM, steering control cells toward drug-treated profiles in a concentration-dependent manner.
6. We release a codebase for training sparse autoencoders on single-cell foundation models that is easily extensible to new autoencoder architectures and foundation models.^1^

## 2. Background

### 2.1. Sparse autoencoders

Sparse autoencoders (SAEs) are neural networks that learn interpretable representations by decomposing neural network activations into sparse, monosemantic features. These latent units respond to single, interpretable concepts rather than multiple unrelated patterns. Recent work has demonstrated their effectiveness for interpreting transformer models, from language models (Cunningham et al., 2023; Bricken et al., 2023; Templeton et al., 2024) to biological sequence models (Guan et al., 2025; Simon & Zou, 2024; Adams et al., 2025; Gujral et al., 2025).

### 2.2. Sparse autoencoders for single-cell foundation models

Schuster (2025) introduced automated interpretability tools for linking SAE features to biological concepts using gene sets, while Claye et al. (2025) developed methods to interpret latent concepts by identifying contributing genes through counterfactual perturbations and leveraging textual gene descriptions from ontologies. Both studies focused primarily on analyzing the cell embedding space of pretrained single-cell models.

Our approach extends this work in several key directions. Unlike previous studies that analyze final cell embeddings, we apply SAEs to intermediate token representations during the model’s forward pass, capturing richer biological information before it is compressed into cell-level summaries. We conduct comparisons across three foundation models (scFoundation, Geneformer, and scGPT under several fine-tuning protocols) and diverse datasets, enabling analysis of how training strategies and data influence learned representations. We also provide a more detailed categorization of discovered features, distinguishing gene- from cell-specific patterns, and introduce novel applications of SAE-based steering to address technical artifacts such as batch effects while preserving biological signals.

## 3. Methods

### 3.1. Sparse Autoencoder Training

We trained sparse autoencoders on intermediate token representations from three single-cell foundation models: scGPT (Cui et al., 2024), scFoundation (Hao et al., 2024), and Geneformer V2 (Theodoris et al., 2023). For each scFM, we performed inference on scRNA-seq datasets and extracted residual stream activations from different transformer layers (Figure 1).

**Figure 1.**
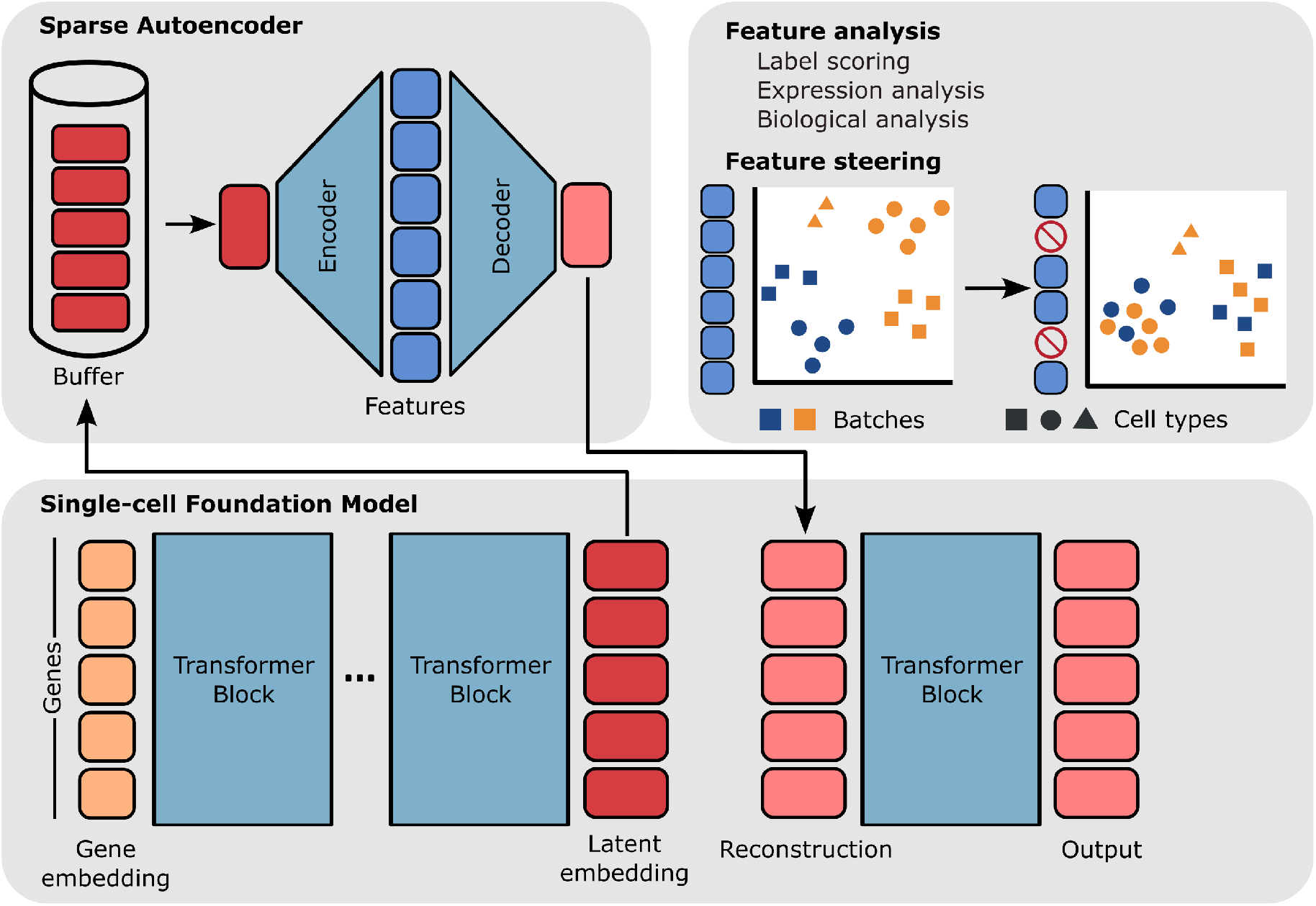
Framework used to train and evaluate sparse autoencoders from scFM latent gene embeddings.

#### Model selection

We analyzed pre-trained and fine-tuned variants where available. scFoundation lacked publicly available fine-tuning code, so we used only the pre-trained model. Pre-trained Geneformer produced poorly-defined features with high reconstruction loss (Figure A.3), so we focused on fine-tuned Geneformer. We prioritize scGPT analysis due to its widespread adoption and fine-tuning ease.

#### SAE architecture

We used BatchTopK SAEs (Bussmann et al., 2024), which outperformed other SAE architectures in both language models and our preliminary experiments (Appendix A.5.1). We trained with Adam (Kingma & Ba, 2014), sampling 250 tokens per cell and constructing batches of 8,192 tokens drawn from varied cellular contexts following Bricken et al. (2023). We kept feature spaces small, as larger dictionaries split concepts into overly fine-grained, uninterpretable features, especially on smaller datasets. Details on the SAE hyperparameters can be found in Appendix A.5.3.

#### Layer selection

We trained SAEs on all 12 transformer layers. Middle layers (5-8) achieved optimal steering performance and captured batch-related features most effectively (Figures A.10, A.12), while later layers (9-11) showed higher reconstruction quality and more structured biological representations (Figures A.2, A.7). We selected layer 6 for steering experiments and layer 10 for interpretability analysis, avoiding final layers which primarily encode prediction-specific features rather than generalizable representations.

#### Evaluation

The standard SAE metrics “loss recovered” (Bricken et al., 2023) failed due to consistently high scFM training loss. To address this limitation, we developed the Embedding Recovery Score, which measures how token-level reconstruction quality affects downstream cell embeddings (Appendix A.5.2).

### 3.2. Feature Analysis

We characterized SAE features through two complementary approaches: cell-level associations and functional enrichment analysis.

#### Cell-level associations

SAEs produce continuous activation values for each feature at each gene position within a cell. We aggregated these gene-level activations to the cell level via max-pooling, yielding a single activation score per feature per cell. This aggregation allows us to ask: does this feature consistently activate for specific cell types, disease states, or technical batches?

To quantify these associations, we thresholded continuous cell-level scores into binary classifications (feature active/inactive) and computed alignment metrics, namely adjusted mutual information (AMI) and F1 scores, against ground-truth cell-level labels. We examined multiple threshold values since the interpretation of “activation” varies and often focused on strongly activated tokens to identify each feature’s core concept. For example, a feature might activate weakly across many B cells but strongly in a specific B cell subtype, revealing finer-grained biological structure.

#### Functional enrichment analysis

To interpret the biological relevance of features, we tested whether genes with strong feature activations were enriched for known biological categories. We performed over-representation analysis (ORA) (Subramanian et al., 2005) using Gene Ontology gene sets (Liberzon et al., 2015) to assess enrichment across biological processes, cellular components, and molecular functions. We also used PanglaoDB marker sets (Franzén et al., 2019) to examine enrichment of cell type-specific expression markers.

Additionally, we computed Spearman’s rank correlations to assess whether features consistently activated for tokens encoding high gene expression values, and quantified feature enrichment for specific gene families by calculating the fraction of each feature’s activations attributed to genes within those families.

### 3.3. Feature Steering

We used steering to test whether identified features contribute to model behavior: if suppressing a feature systematically alters model outputs in predictable ways, this provides evidence that the feature actively encodes that property rather than representing spurious correlation. Specifically, we tested whether SAE features can be manipulated to reduce batch effects while preserving the biological signal.

#### Suppression (batch effect correction)

We identified batch-correlated features by computing adjusted mutual information (AMI) between feature activations and batch labels (Section 3.2), and clamped these “batch features” to −2 whenever they activated, suppressing the technical signal. We explored different clamp values (Figure A.11) and −2 balanced batch correction against biological signal preservation. We clamped 50 features for fine-tuned models and 20 for pre-trained models, where batch-correlated features were less distinct.

#### Induction (drug perturbation response prediction)

Conversely, we tested whether activating a feature could induce a perturbation-associated state. We identified features tracking drug treatment and concentration (Section 3.2) and steered control cells toward treated states by clamping selected drug-perturbation features to a high value (its maximum observed activation) in a randomly chosen fraction of each cell’s gene tokens. Varying this fraction results in gradual steering. Unlike suppression, which is applied at every position where a feature fires, induction is applied to a token subset, with the fraction controlling steering strength.

#### Evaluation

For batch suppression, we compared steered embeddings against: (1) original scFM embeddings, (2) PCA on raw gene counts (no batch removal), and (3) scVI (Lopez et al., 2018), a conditional VAE-based batch correction method. Following Luecken et al. (2022), we evaluate batch correction along two axes: bio conservation and batch mixing. Bio conservation measures how well the learned representation preserves the biological signal of interest, in our case cell type labels, while batch mixing measures how well cells from different batches are intermixed. The metrics are further detailed in Appendix A.2. scVI serves as a reference point for batch effect magnitude rather than a competitive baseline. For drug induction, we assessed whether steered control cells moved toward the perturbed population using two metrics: the Cluster Merging Score, the fraction of a steered cell’s k nearest neighbors that are perturbed rather than control cells; and the Directional Alignment Score, the normalized projection of each embedding onto the control-to-perturbed centroid axis (Appendix A.3).

### 3.4. Datasets

We trained SAEs on six datasets: CellXGene Census (37M cells) (CZI Cell Science Program et al., 2025), a COVID-19 cohort (Yoshida et al., 2022), a subset of the Tahoe-100M drug perturbation dataset (Zhang et al., 2025), and three tissue-specific datasets (immune, lung, pancreas) from Luecken et al. (2022). These three datasets were compiled for a batch integration benchmarking study and provide controlled scenarios with varying batch effects and cell type compositions that can be used for steering validation. COVID-19 and CellXGene Census enabled large-scale feature characterization. Details in Appendix A.1.

## 4. Results

### 4.1. Sparse Autoencoders find interpretable concepts

Sparse autoencoders trained on scGPT, scFoundation, and Geneformer revealed that scFMs organize information along two distinct axes: gene-specific features that encode properties of individual genes independent of cellular context, and cell-specific features that capture properties of entire cells, distributed across multiple gene tokens. This decomposition reveals how transformer architectures integrate local (gene) and global (cell) information during sequence processing.

#### Gene-specific features encode expression levels, gene identity, and molecular function

These features activate based on properties intrinsic to individual gene tokens. Across all three models, we identified features correlating with gene expression values, though the encoding strategy varied by architecture (Tables A.3, A.2, Figure A.4). scFoundation features primarily captured low expression values, likely because this model applies Bayesian downsampling during training, making high expression values inconsistent indicators of relative abundance. scGPT showed stronger expression correlations across multiple expression levels due to its binning strategy, which maps expression ranges to discrete bins and removes technology-based variability. Geneformer, which encodes expression through token position rather than token value, produced features that activated most often at specific sequence positions.

Beyond expression, gene-specific features captured gene identity and family membership. We found features selective for ribosomal, mitochondrial, HLA, and immunoglobulin gene families (Tables A.4, A.5, A.6). A third category encoded biological processes such as cell cycle, defense response and apoptosis through activation on functionally related gene sets (Tables A.7, A.8, A.9).

#### Cell-specific features encode cell identity through distributed representations

Unlike gene-specific features, these activate based on the broader cellular context in which a gene appears. They arise from contextual information that transformers propagate across the sequence during self-attention, often distributed across many tokens rather than concentrated in the most biologically relevant genes.

In the COVID-19 dataset, we identified features corresponding to major cell types across all models, with broader categories (e.g. monocytes, B cells, T cells) represented more consistently than fine-grained subtypes (Tables A.10, A.11, A.12). These cell type features were often enriched for appropriate marker genes and biological processes. However, feature structure varied across models, with scGPT having substantially more numerous and diverse feature types per cell type than scFoundation or Geneformer (Figure 2). These cell type features could be categorized as marker gene selective, generalized, ribosomal, or low expression based on their activation patterns. scGPT produced 3-8 diverse features per cell type even before fine-tuning, while scFoundation most often generated 2-3 complementary features per cell type. Geneformer required fine-tuning to develop clear representations and showed predominantly generalized features.

**Figure 2.**
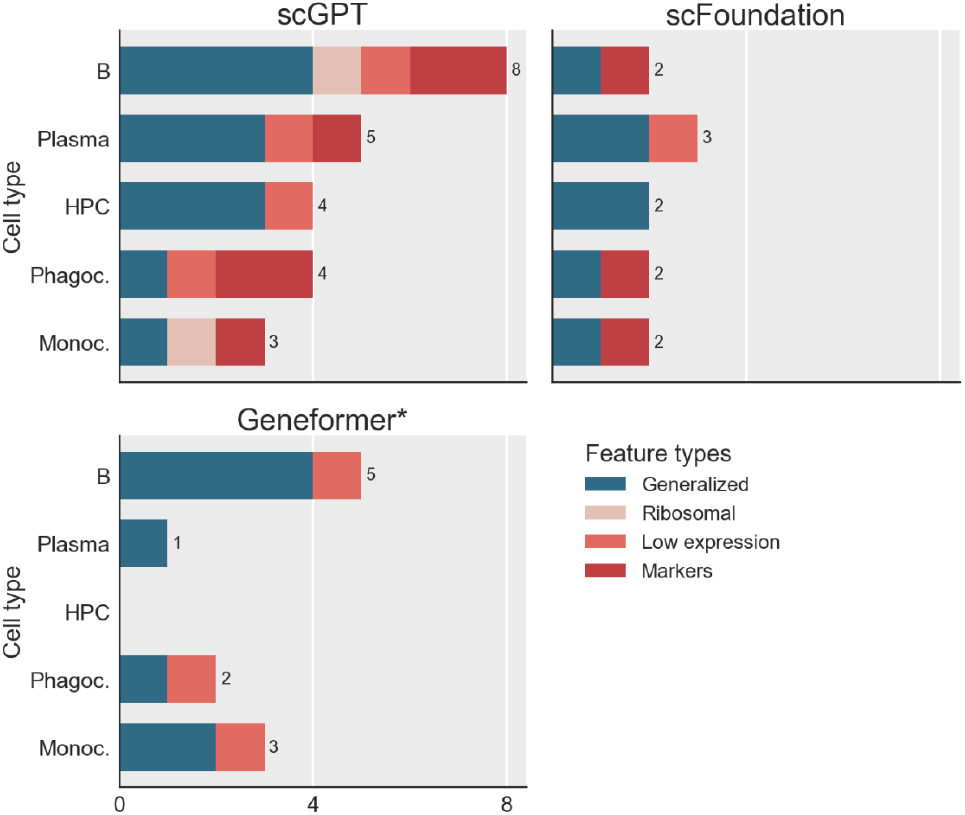
Distribution of feature types across the best represented cell types in the scFMs. Stacked bars show the number of features identified for each cell type, categorized by activation pattern: generalized features that activate broadly across genes and expression levels, marker features selective for canonical cell type markers, ribosomal features activating on ribosomal genes, and low expression features that preferentially activate on lowly expressed genes. Numbers indicate total features per cell type. *Geneformer needed to be fine-tuned before showing cell type-specific features while the other two models are pre-trained here.

Beyond cell types, we found features encoding disease status (activating preferentially in COVID-19 patients) and technical batch effects (activating for specific studies, donors, or sequencing technologies) (Tables A.13, A.14, A.15, A.16).

### 4.2. Concepts from pre-trained scFM are varied and meaningful

The following findings demonstrate that pre-trained scFMs can develop rich internal representations even before task-specific fine-tuning. These representations are compositional, combining multiple distinct features to encode cell identity and transfer to contexts not present during training, such as disease states. We focus on scGPT, as it best exemplified these properties among the pre-trained models we analyzed.

#### Features capture diverse aspects of cell identity

For each cell type, we observed several distinct features with different characteristics, suggesting scFMs build cell representations from compositional features rather than unified cell type detectors. Table 1 illustrates this using B-cell features from pre-trained scGPT.

**Table 1.** Selected scGPT features predictive of B cell identity with key characteristics. Expr reports mean ± SD expression. # Genes denotes the number of distinct genes activated by the feature. Ribo (%) indicates the percentage of active genes that are ribosomal.

| Feat | AMI | F1 | Expr | # Genes | Ribo (%) |
| --- | --- | --- | --- | --- | --- |
| 478 | 0.97 | 0.99 | $18.7 \pm 6.1$ | 1485 | 0 |
| 363 | 0.95 | 0.99 | $42.6 \pm 5.8$ | 83 | 0 |
| 151 | 0.99 | 1.00 | $4.3 \pm 2.9$ | 2698 | 2 |
| 270 | 0.85 | 0.95 | $43.9 \pm 5.4$ | 187 | 85 |

Some features activated broadly across many genes and expression levels, representing general cell type membership (e.g., feature 478). Other features targeted specific marker genes: feature 363 activated strongly (activation ≥ 0.5) on only 83 genes, 26 of these were known B-cell markers and another 38 immunoglobulin or MHC genes (both commonly enriched in B cells). Gene Ontology analysis confirmed enrichment for B-cell-mediated immunity (adjusted p-value = 2.1e-17). This feature thus captures a highly specific B-cell signature through canonical marker gene activity. While these two features follow expected patterns by activating on canonical markers or broadly across B-cell contexts, we identified two additional unexpected encoding strategies.

First, feature 151 implements *negative encoding*: it activates moderately on genes enriched for T-cell, monocyte, macrophage, megakaryocyte, and NK cell markers, but notably not B-cell markers. Rather than directly detecting B-cell signatures, this feature encodes “absence of non-B-cell signatures” within B cells, effectively serving as a negative indicator for other cell identities.

Second, feature 270 demonstrates *proxy encoding*, activating on specific ribosomal gene subgroups whose expression levels differentiate B cells from other cell types despite lacking direct biological association with B-cell function. This suggests the model exploits any discriminative pattern to construct cell representations, whether or not it aligns with canonical biological markers.

#### Features encode biological processes from unseen conditions

Pre-trained scFM models can capture distinct aspects of cellular function and state, including features that reflect specific biological processes or abnormal cell states not present in the training data.

Despite training only on healthy cells, scGPT developed features that activated in disease-associated cellular states. One feature activated predominantly in monocytes and dendritic cells from patients with post-COVID-19 disorder (Figure A.5), with strong enrichment for inflammation-related pathways (adjusted p-value: 1.19e-23). This aligns with clinical observations that monocytes and dendritic cells in post-COVID patients remain in persistently activated inflammatory states months after acute infection (Boes & Falter-Braun, 2023; Hopkins et al., 2023).

#### Features capture technical aspects of sequencing protocols

Pre-trained scFM models also capture systematic biases inherent in sequencing technologies. These biases manifested as features that correlate with technical variables such as gene length and GC content.

In the raw gene expression values from the pancreas dataset, we observed that cells processed with the sequencing protocol “SMARTer” showed distinct technical signatures compared to other sequencing methods. Specifically, these cells exhibited a stronger positive correlation between gene count and gene length, and a stronger negative correlation with GC content, compared to the dataset as a whole. The scGPT feature with one of the highest AMI scores between its activations and the SMARTer sequencing label captured exactly this technical signature: it activated on genes with significantly greater length and lower GC content than the dataset average (Figure 3), specifically when these genes were highly expressed.

**Figure 3.**
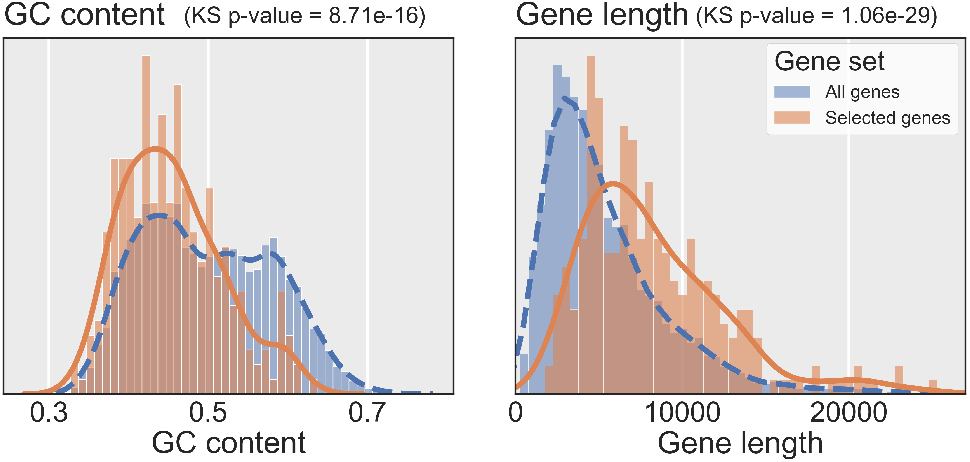
Density histogram of gene GC content and length distribution for all genes in the dataset (blue) and those with highest activation in feature 236 (orange).The SAE was trained on the scGPT model using the pancreas dataset. KS refers to the Kolmogorov-Smirnov test statistic.

### 4.3. Features capture technical variation alongside cell type information

The CellXGene Census dataset aggregates many studies using different experimental protocols and sequencing technologies. A key requirement for single-cell foundation models is their ability to generalize beyond protocol-specific technical artifacts and capture shared biological principles. While technical effects persist in learned representations unless explicitly addressed during training, robust models should identify biological concepts that extend across these experimental variations.

To investigate how pre-training on this dataset shapes scGPT’s internal representations, we trained a sparse autoencoder on the model’s intermediate tokens. Our cell-level association analysis revealed that many features correlated with specific datasets, while others showed weaker correlations with sequencing technologies. Representative examples of these features are provided in Table A.17. This finding, though unsurprising, demonstrates substantial interdataset variability and suggests that the model may allocate considerable representational capacity to encoding these technical differences.

Many features also showed varying generalization across studies and technologies. While some demonstrated consistent activation patterns census-wide, many associated more strongly with specific data subsets. SAE-derived cell type concepts showed variable activation, with some activating on cells from particular studies while showing reduced activity on the same cell types from other studies.

To quantify this, we compared the AMI between feature activations and complete cell type annotations versus cell type subsets spanning study subgroups. Study subsets typically showed stronger alignment than complete cell type categories (Figure A.6), suggesting scGPT captures cellular identity across some studies but may not consolidate signals into unified concepts spanning all instances census-wide.

### 4.4. Cell embeddings can be integrated by deactivating specific features

Having identified that SAE features encode batch effects (Section 4.3), we tested whether these features actively contribute to technical variation in model outputs.

We identified the top batch-correlated features (by AMI) in scGPT, Geneformer, and scFoundation, and clamped them to −2 during inference (see Methods 3.3). We clamped 50 features for fine-tuned models but only 20 features for pre-trained models, as batch-correlated features were less distinct in these models. Table 2 shows results for the pancreas dataset; Tables A.19 and A.20(Appendix) report lung and immune results.

**Table 2.** Batch correction performance of single-cell foundation models on the pancreas dataset. Values show batch correction, biological conservation, and total scores, with higher scores indicating better performance. Means and standard deviations are computed across five runs with different random model initializations. Bold values highlight the best method within each category.

| Method | Batch correction | Bio conservation | Total |
| --- | --- | --- | --- |
| Unintegrated (PCA) | 0.11±0.000 | 0.54±0.015 | 0.37±0.009 |
| scVI | 0.85±0.027 | 0.81±0.010 | **0.82**±0.014 |
| scFoundation (zero-shot) | 0.32±0.000 | 0.36±0.010 | 0.34±0.006 |
| scFoundation (zero-shot, steered) | 0.47±0.000 | 0.29±0.001 | **0.36**±0.006 |
| Geneformer (fine-tuned) | 0.56±0.000 | 0.86±0.007 | 0.74±0.005 |
| Geneformer (fine-tuned, steered) | 0.65±0.00 | 0.87±0.005 | **0.78**±0.003 |
| scGPT (zero-shot) | 0.60±0.016 | 0.04±0.011 | 0.26±0.010 |
| scGPT (zero-shot, steered) | 0.60±0.029 | 0.07±0.013 | **0.31**±0.014 |
| scGPT (fine-tuned) | 0.42±0.007 | 0.47±0.005 | 0.45±0.005 |
| scGPT (fine-tuned, DAR) | 0.58±0.034 | 0.58±0.007 | 0.58±0.012 |
| scGPT (fine-tuned, steered) | 0.70±0.038 | 0.56±0.005 | **0.62**±0.013 |

As expected, unintegrated PCA embeddings achieved the lowest batch-correction score, since no batch removal is applied, while the explicit batch integration method scVI achieved the highest total score. Steering improved batch correction for fine-tuned models across all datasets without substantial loss in biological conservation. For the pancreas dataset, steering the fine-tuned scGPT outperformed both the native Domain Adaptive Regularization (DAR) batch correction of scGPT and the unsteered fine-tuned model, with improved biological conservation. Appendix Figure A.8 shows Uniform Manifold Approximation and Projections (UMAPs) (Healy & McInnes, 2024) of embeddings from unaltered and steered fine-tuned scGPT, colored by cell type and sequencing technology. These plots show how steering reduces batch effects and improves clustering by cell type. Pre-trained steering was only consistently successful on the pancreas dataset. This likely reflects the stronger batch effects in this dataset and the fact that batches correspond to sequencing technologies the models encountered during pre-training, rather than donor-or laboratory-specific effects.

Pre-trained scGPT embeddings showed lower batch effects than those of the model fine-tuned on the specific dataset. Fine-tuning without correction caused the model to internalize batch effects, since doing so improved its ability to minimize the training loss for gene expression prediction.

Finally, we compared the effects of deactivating different numbers of features using steering versus randomly selected features (Figures 4 and A.9). For the pancreas dataset, performance increased for up to 25 deactivated features and then plateaued. In contrast, for the lung and the immune datasets, performance increased for up to 30 and 60 deactivated features, respectively, but then decreased significantly beyond these thresholds. Furthermore, we evaluated different clamping values in Appendix Figure A.11. The experiment showed that preservation of biological signal decreases with lower clamping values, while batch correction and total score reach their highest values for −2.

**Figure 4.**
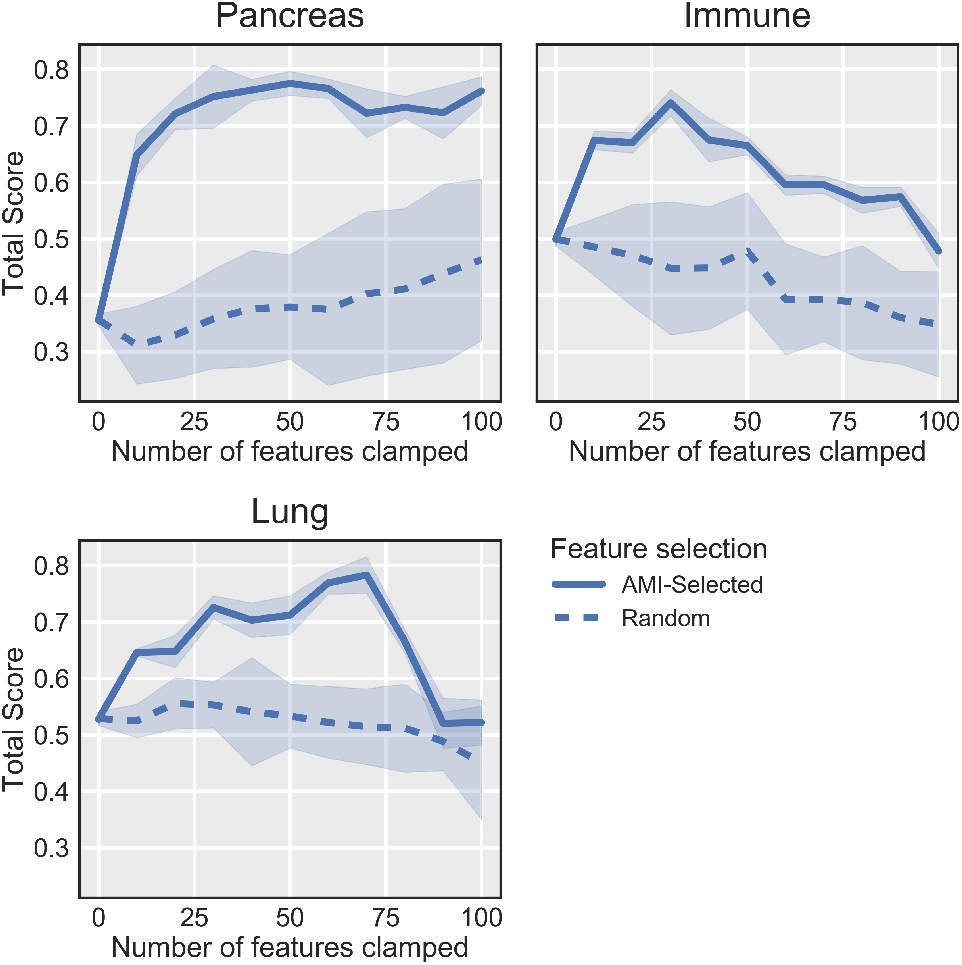
Total integration score as features are sequentially steered, selected randomly or by maximum AMI, across three datasets. Lines represent the mean across five seeds, and shaded regions indicate standard deviation.

### 4.5. Activating features induces drug-perturbation states

The steering experiments above suppress unwanted signals. We next asked whether the inverse operation, activating a feature, could induce a desired biological state. We tested this on a subset of Tahoe-100M (Zhang et al., 2025) comprising 924,896 cells treated with three concentrations for three different drugs (Dinaciclib, Panobinostat and Bortezomib). We trained the SAE on all cell lines except the three largest per drug (six total due to overlap), which we held out for evaluation, testing whether dose-encoding features generalize to cell lines unseen during SAE training. To ensure that perturbation information was available to the SAE, the underlying scFMs were first fine-tuned on the same data.

We identified features whose activations tracked drug treatment (Section 3.2) and used them to steer control cells toward treated states. Rather than clamping features to a negative value at every firing position, as in batch suppression, we clamped a selected dose-encoding feature to a high value in a random subset of each cell’s gene tokens (Section 3.3). We then measured whether steered control cells moved into the perturbed population using the Cluster Merging Score (CMS) and the Directional Alignment Score (DAS) (Appendix A.3).

Activating dose-encoding features shifted control cells toward the drug-treated population across all three drugs. Figure 5 shows embeddings of control and treated cells before and after steering: steered control cells move out of the control cluster and intermix with treated cells. After steering, the mean CMS across cells exceeded 0.9 for all three drugs, indicating that steered cells are surrounded almost entirely by perturbed neighbors (Figure 6). The DAS ranged from approximately 0.7 to 1.05, placing steered cells near or slightly beyond the perturbed centroid and confirming substantial displacement along the control-to-perturbed axis. Because evaluation cell lines were absent from SAE training, this shift reflects a dose feature that generalizes beyond the training distribution rather than memorization of specific cell-line profiles.

**Figure 5.**
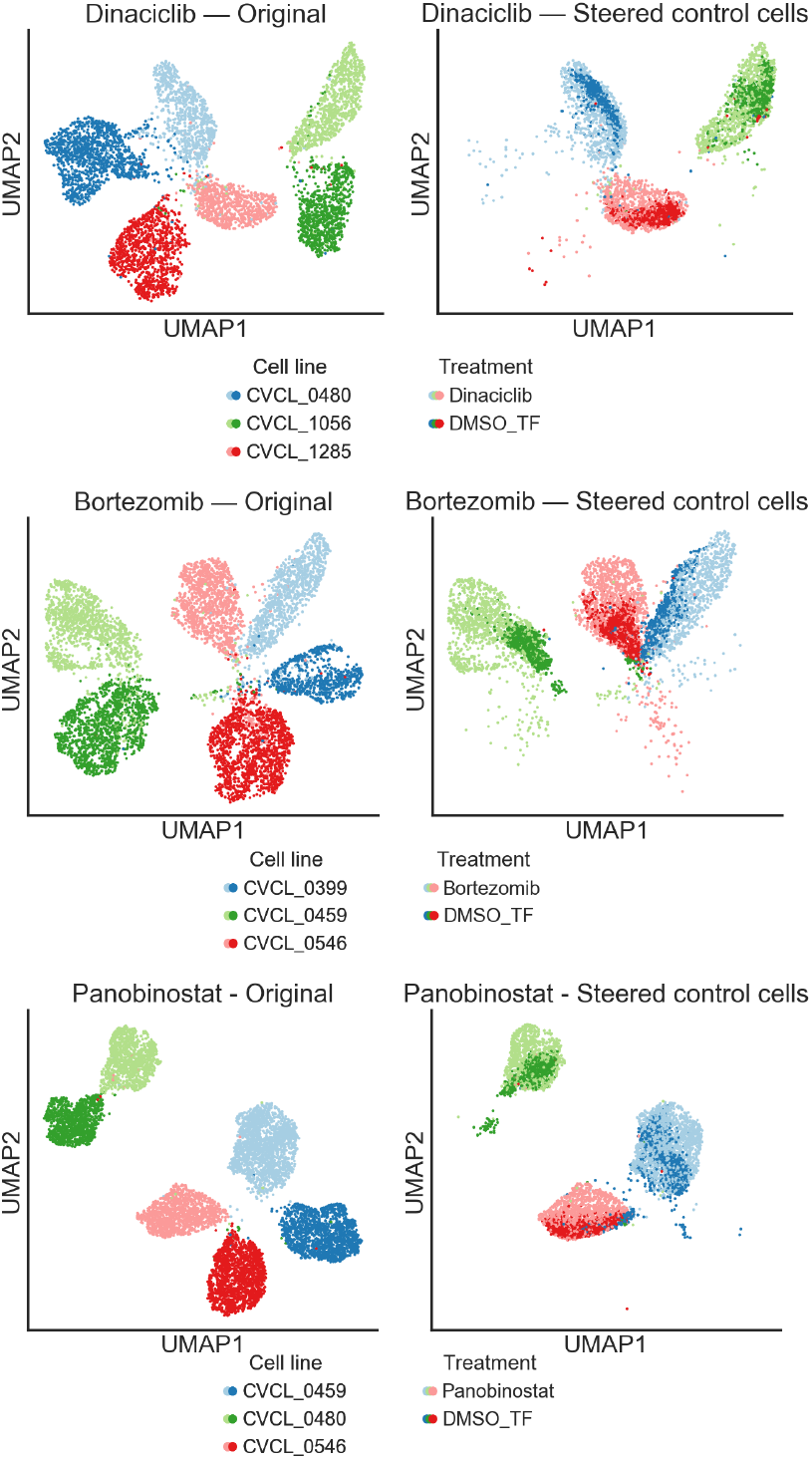
UMAPs of scGPT embeddings for cells treated with Bortezomib (top), Dinaciclib (middle), and Panobinostat (bottom). Each panel shows original embeddings (left) and a mixed representation in which control (DMSO) cells are replaced by their SAE-steered counterparts (right). Colors encode cell line and treatment condition; darker shades denote drug-treated cells, lighter shades denote controls.

**Figure 6.**
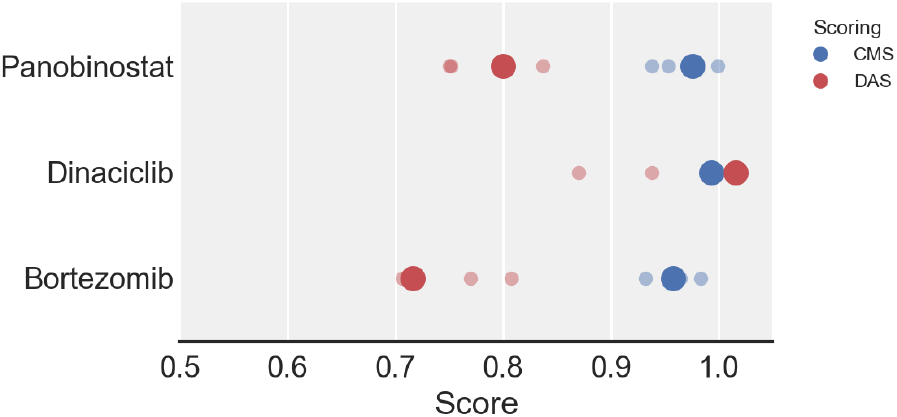
Cluster Merging Score (CMS) and Directional Alignment Score (DAS) for each steered perturbation effect. Large dots indicate the pooled score across all control cells; small dots indicate per-cell-line scores.

Beyond the binary treated-versus-control shift, we examined whether steering strength could track drug concentration. For Panobinostat, whose treated embeddings form a concentration-graded continuum, increasing the fraction of clamped tokens moved steered controls progressively along the control-to-perturbed axis, with their DAS distributions sweeping through those of successively higher real concentrations (Figures 7 and A.13). These results suggest that feature activation can induce perturbation-associated states, and that where the underlying representation is dose-graded, steering magnitude reproduces that grade.

**Figure 7.**
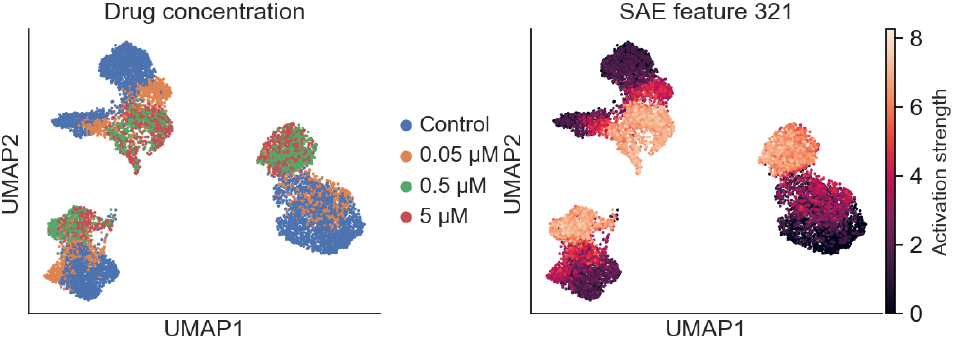
UMAPs of scGPT embeddings of Panobinostat treated and control cells coloured by drug concentration (left) and by the activation strength of SAE feature 321 (right), computed as the maximum activation across all input tokens.

## 5. Discussion & Conclusion

### scFMs learn meaningful biological representations during pre-training

Decomposing latent representations through SAEs reveals that scFMs develop a nuanced understanding of cell biology during pre-training.

The gene-specific versus cell-specific decomposition we observed reflects how transformers process sequential biological data. Gene-specific features operate locally, capturing token-level properties like expression values and gene family membership, while cell-specific features emerge from global context aggregation across self-attention layers. This distinction has implications for model interpretability: gene features directly reflect biological mechanisms (gene function, expression regulation), while cell features reveal how models synthesize distributed evidence across many tokens to infer cellular identity and often through unexpected strategies like negative encoding or proxy markers rather than canonical biological signatures alone.

While current studies report that scFMs underperform in strict zero-shot settings, the presence of rich biological features suggests that continued model development may close this gap and lead to foundation models with stronger zero-shot performance. However, a substantial proportion of features remains difficult to characterize. Standard methods such as Gene Ontology analysis prove insufficient, as many features reflect heuristic gene co-expression patterns whose biological relevance is challenging to establish. This interpretability gap is larger than in language models, where features often map to human-interpretable concepts.

### Training procedures affect information encoding

Our cross-model comparison revealed systematic differences in how scFMs represent biological information such as gene expression encoding and cell type feature diversity.

Expression encoding significantly impacts feature structure, with models adopting different strategies to deal with technology-based variance in gene expression. scGPT and Geneformer attempt to remove expression variance through binning and ranking strategies while scFoundation is explicitly trained to be robust to these fluctuations. These different strategies manifest clearly in SAE features. scGPT’s binning produced features with strong expression correlations across multiple expression levels, as consistent bin mappings allow the model to reliably identify expression patterns. Geneformer’s position-based encoding yielded features that activated preferentially at specific sequence positions rather than for expression magnitudes. scFoundation showed fewer features strongly correlated with high expression values, likely because downsampling makes high expression an inconsistent indicator of relative abundance, training the model to attend to other signals.

scGPT also generated more diverse features per cell type than other models, even before fine-tuning, with notably more ribosomal-focused features. This may relate to its pre-training approach: higher masking ratios (25-75% vs. 15% in Geneformer and 30% in scFoundation) and shorter gene sequences force the model to infer cell context from broader gene sets, potentially requiring more diverse feature repertoires.

These findings suggest that architectural choices can have strong effects on learned representations, warranting systematic comparison to determine which strategy better captures biological signal while minimizing technical artifacts.

#### Representations reflect technical variation but exhibit limited cross-study consolidation

Our analysis revealed that scGPT dedicates considerable representational capacity to encoding technical differences between datasets, with many features correlating strongly with specific studies or sequencing technologies. This highlights a fundamental challenge in label-free pre-training: models will learn to represent strong technical signals regardless of biological relevance.

The variable generalization of cell type features across studies could partly reflect SAE limitations in how representations are decomposed, but may also indicate genuine fragmentation in underlying model representations. Because pre-training is label-free, models learn to distinguish cells based on observed gene expression patterns without knowing which patterns reflect biology versus technical variation. When technical factors such as sequencing protocols, laboratory conditions, or sample preparation methods create strong signals that overshadow shared biological patterns, models may learn separate representations for the same cell type across studies, treating them as distinct entities. Fine-tuning may be essential not because it adds new information, but because it recontextualizes existing information, linking study-specific representations and allowing models to recognize that separately learned representations are instances of the same biological concept.

### SAE features are functionally related to model behavior

Steering experiments demonstrate that high-AMI features actively encode correlated information rather than spurious associations. Suppressing batch-related features systematically improved batch integration metrics, confirming these features functionally contribute to technical variation. Steering also operates in the opposite direction: activating dose-encoding features induced drug-perturbation states, moving control cells into the treated population across three drugs while recreating the concentration spectrum. That the same intervention both removes unwanted signals and induces desired ones, and that induced shifts generalize to held-out cell lines, indicates SAEs capture functionally relevant and manipulable directions in scFMs rather than descriptive correlations alone.

These results also show model representations are decomposable: intervening on specific features selectively alters behavior without disrupting other tasks. This modularity could aid understanding and controlling model behavior in single-cell applications. While current steering isn’t robust enough for batch correction, it opens possibilities for sophisticated interventions. Batch information could be explicitly encoded during training for targeted removal. The ability to identify and manipulate features suggests applications in removing biases or technical artifacts from pre-trained models, and conversely in inducing states such as perturbation responses, analogous to concept editing in large language models.

### Conclusion

We applied sparse autoencoders to reveal the internal mechanisms of single-cell foundation models, finding that these models develop compositional, generalizable representations during pre-training but fragment them across study-specific contexts. Our cross-model comparison demonstrated how architectural choices affect representation structure, while steering experiments confirmed that learned features causally encode their associated properties and can be manipulated to reduce technical artifacts or induce drug perturbations. While the method shows potential, applying SAEs in the scRNA-seq context proves substantially more challenging than in language models, with feature interpretation requiring extensive manual effort and automated approaches showing limited success.

As single-cell foundation models remain in early development without consolidated best practices, interpretability studies examining what information these models encode and how training protocols affect representations will be essential for moving toward more reliable and controllable models. The tools and findings presented here provide a foundation for such investigations.

## A. Appendix

### A.1 Datasets

All benchmark datasets utilized in our study are openly accessible to the public.

#### CellXGene Census

The Census provides an efficient tool to access and query all single-cell RNA data from CZ CELLxGENE Discover (CZI Cell Science Program et al., 2025). By querying for healthy cells across a range of tissues, we obtained a dataset of over 37 million cells. This dataset was reportedly used by the authors of scGPT for model training.

#### COVID-19

The COVID-19 dataset (Yoshida et al., 2022) includes 33,105 genes measured in 422,220 peripheral blood mononuclear cells, covering 16 annotated cell types from both healthy individuals and COVID-19 patients. Availability: https://datasets.cellxgene.cziscience.com/ae49598b-646d-4325-b3e7-b164ac49d506.h5ad

#### Immune

This collection consists of 33,506 cells containing 12,303 genes sourced from ten distinct donors, compiled by Luecken et al.(2022) across five research studies. While one investigation obtained cells from human bone marrow, the remaining four studies extracted cells from human peripheral blood. The dataset contains annotations for 16 distinct cell types. Availability: https://doi.org/10.6084/m9.figshare.12420968.v8

#### Pancreas

This collection was reprocessed by Luecken et al.(2022) through the integration of five human pancreas studies. It encompasses 16,382 cells, featuring 19,093 genes, sequenced using four scRNA-seq platforms (inDrop, CEL-Seq, Smart-Seq2, SMARTer). The integrated dataset incorporates 14 cell types. Availability: https://figshare.com/ndownloader/files/24539828

#### Lung

This collection encompasses 32,426 cells spanning 16 batches and two sequencing platforms (Drop-seq and 10x Chromium), compiled by Luecken et al.(2022) from three research laboratories. The integrated dataset incorporates 15,148 genes. The cells originate from transplant patients and lung tissue samples and are classified into 17 cell types. Availability: https://figshare.com/ndownloader/files/24539942

#### Tahoe-100M (subset)

We selected a subset of Tahoe-100M (Zhang et al., 2025) comprising 924,896 cells treated with three drugs (Dinaciclib, Panobinostat, Bortezomib) across 3 concentrations. We trained the SAE on all cell lines except the six largest, which we held out for steering evaluation, providing a test of whether dose-encoding features generalize to cell lines unseen during SAE training.

### A.2. Batch correction metrics

To evaluate how well different methods integrate cells from different experimental batches, we follow (Luecken et al., 2022). The evaluation metrics are organized into two distinct groups: one set focuses on assessing how well the biological variability is preserved, while the other assesses the effectiveness of aligning cells from different batches.

For assessing the preservation of biological variation, we employ several metrics, including the isolated labels score, normalized mutual information (NMI) and adjusted rand index (ARI), silhouette label score, and the cLISI metric. For measuring batch correction performance, we utilize graph connectivity analysis, kBET calculations per label, individual cell iLISI values, PCR comparison scores, and batch-specific silhouette coefficients.

The bio conservation and batch correction scores are computed by first min-max normalizing each individual metric and then taking the mean across all bio conservation or batch correction metrics, respectively. The total score is calculated as 0.6× bio conservation + 0.4× batch correction. Comprehensive descriptions of these metrics can be found in (Luecken et al., 2022).

### A.3. Drug perturbation metric

We quantify drug-perturbation steering with two complementary metrics, both computed in the cell embedding space. Let 𝒫 and 𝒞 denote the sets of genuinely perturbed and control cell embeddings, respectively, and let *x* be a steered control cell embedding.

#### Cluster Merging Score (CMS)

The CMS measures whether a steered cell has moved into the perturbed population by inspecting the composition of its local neighborhood. For a steered cell *x*, let 𝒩_*k*_(*x*) be its *k* nearest neighbors (by Euclidean distance) among the combined set of perturbed and control cells. The score is the fraction of those neighbors that are perturbed:

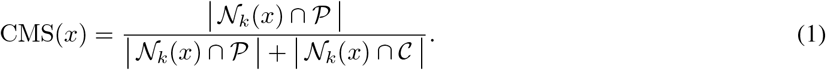

A value near 1 indicates the steered cell is surrounded by perturbed cells (successful merging), while a value near 0 indicates it remains among controls. We report the mean CMS over all steered cells and within cell lines. The CMS treats perturbation as a binary distinction between treated and control populations and is therefore applied to the maximum-concentration steering setting, where all treated cells form the perturbed class.

##### Directional Alignment Score (DAS)

The DAS measures the magnitude of the induced shift along the axis separating control and perturbed populations, capturing graded displacement rather than discrete cluster membership. We define the control-to-perturbed axis as the difference between population centroids,

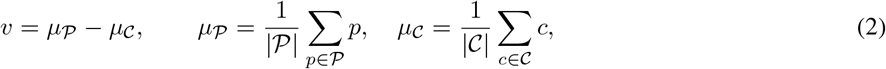

and project each embedding onto this axis, normalizing so that the control centroid maps to 0 and the perturbed centroid to 1:

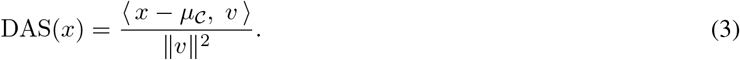

Under this normalization, DAS(*x*) = 0 for a cell at the control centroid and DAS(*x*) = 1 for a cell at the perturbed centroid; intermediate values indicate partial displacement, and values outside [0, 1] indicate cells positioned beyond either centroid along the axis. Because the DAS is continuous, it is suited to the dose-gradient analysis where treated cells do not form discrete clusters.

### A.4. scFM fine-tuning protocols

In all experiments scGPT fine-tuning refers to continued training of the pre-trained model on a new dataset using only the self-supervised objectives GEP (Gene Expression Prediction) and GEPC (GEP for Cell Modelling) as described in the original scGPT paper, allowing the model to learn the target dataset’s distribution without specific label-based tasks. The DAR (Domain Adaptive Regularization) objective was additionally used when explicitly stated.

Geneformer on the other hand, was fine-tuned using supervised learning with cell type labels. However, we removed the CLS token from the latent token embeddings before analysis.

### A.5. Sparse Autoencoders

#### A.5.1. Architectures

Sparse autoencoders are a relatively new approach to discovering interpretable features in foundation models. Several methods exist for applying a sparsity penalty to the feature space. The early version by Bricken et al. (2023) simply applies an L1 regularization penalty to the feature activations. Gated sparse autoencoders (Rajamanoharan et al., 2024) decouple the detection of which features are active from the estimation of their magnitudes. BatchTopK sparse autoencoders (Bussmann et al., 2024) use an activation function that retains only the k largest latents per batch, discarding the L1 regularization entirely. Matryoshka sparse autoencoders (Bussmann et al., 2025) do not address the sparsity penalty, but instead introduce a hierarchical feature structure by training multiple nested dictionaries of increasing size. Smaller dictionaries are forced to independently reconstruct the inputs without relying on the larger ones. The aim is to reduce the incentive created by the sparsity penalty for more specific concepts to absorb high-level features (Chanin et al., 2024).

#### A.5.2. Embedding Recovery Score

The Embedding Recovery Score estimates the impact of token-level reconstruction on downstream embeddings by comparing cell embeddings produced by the model under different conditions:

- **h**_original_: embedding from the original input,
- **h**_reconstructed_: embedding after reconstructing the tokens,
- **h**_0_: ablated embedding where all tokens are zeroed out.

**Figure A.1.**
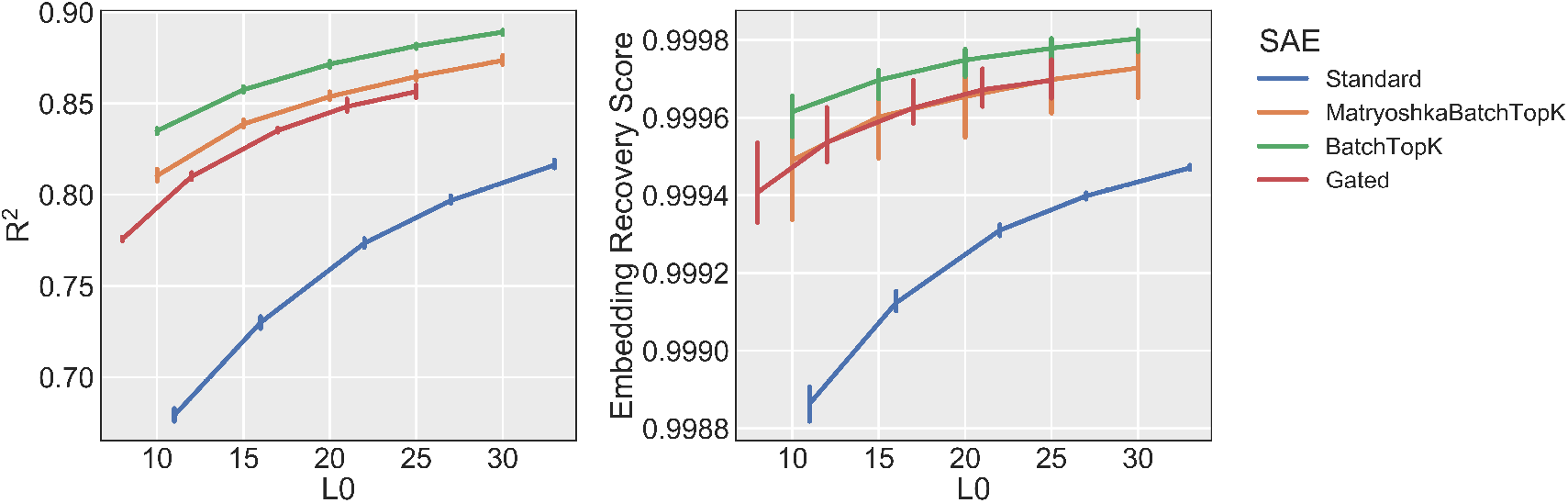
Performance of the four different SAE architectures at different sparsity levels trained on scGPT and the Covid-19 dataset.

The score is defined as

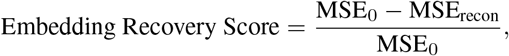

with

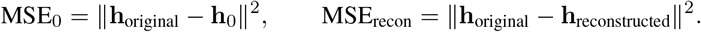

The Embedding Recovery Score inherits some limitations from the loss recovered metric. Notably, zero-ablation has been criticized as an overly pessimistic baseline for defining the zero-point of this metric (Rajamanoharan et al., 2024). While we observed some instability in this metric during early training stages, it remained sufficiently stable for our analysis purposes.

#### A.5.3. Training Hyperparameters

All sparse autoencoders were trained by sampling residual stream activations without replacement from the scFM with a batch size of 8,192. The SAE architecture used for all experiments was a one hidden layer MLP with BatchTopK sparsity restriction. The SAE architecture and trainers were adapted from (Marks et al., 2024).

Hyperparameters were adjusted according to dataset size and complexity as seen in Table A.1. Smaller datasets (Pancreas, Lung, Immune) used a higher learning rate of 1e-3, while larger datasets (COVID-19, CellxGene) used a lower learning rate of 1e-4 for more stable training. The CellxGene dataset used a larger latent dimension (1024 vs 512) and higher sparsity value (20 vs 10) to accommodate its increased variability and provide better expressiveness. The sparsity term refers to the top k active neurons per batch as described by Bussmann et al. (2024).

**Table A.1.** Sparse autoencoder hyperparameters for each dataset.

| Dataset | Learning Rate | Latent Dimension | Sparsity |
| --- | --- | --- | --- |
| Pancreas | 0.001 | 512 | 10 |
| Lung | 0.001 | 512 | 10 |
| Immune | 0.001 | 512 | 10 |
| COVID-19 | 0.0001 | 512 | 10 |
| CellxGene | 0.0001 | 1024 | 20 |

#### A.5.4. Reconstruction losses

**Figure A.2.**
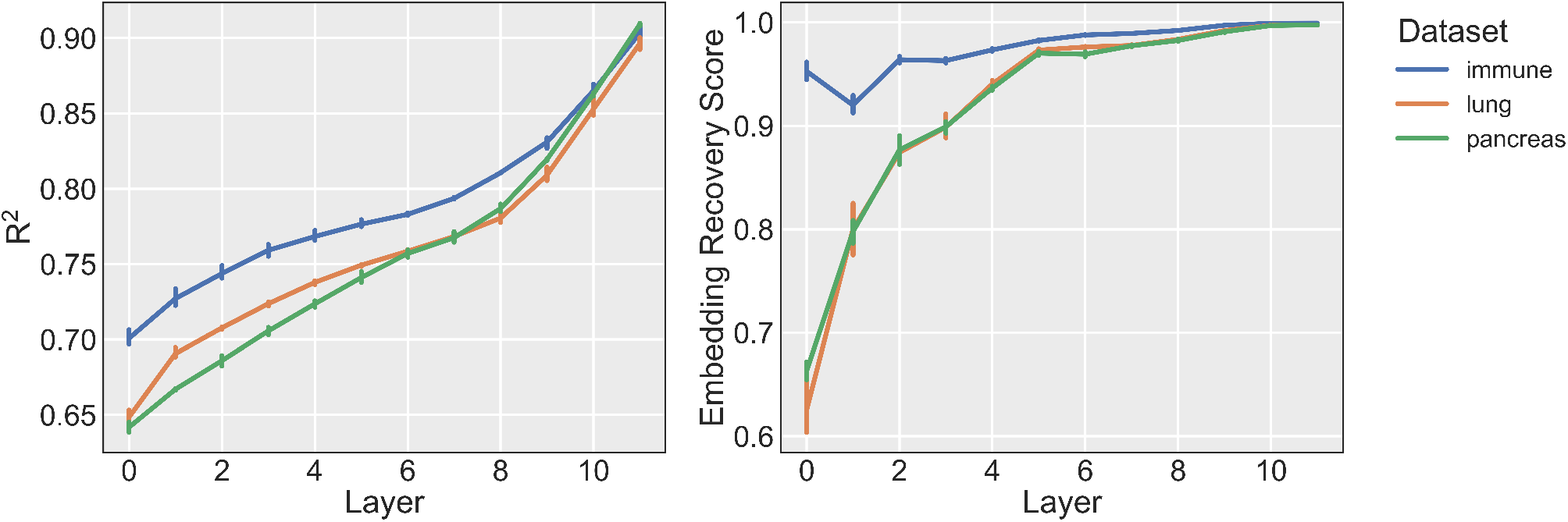
The final training metrics *R*^2^ and Embedding Recovery Score for SAEs trained on different layers of scGPT across three datasets: Lung (blue), Immune (orange), and Pancreas (green). Lines represent the mean across three seeds, and bars indicate standard deviation.

**Figure A.3.**
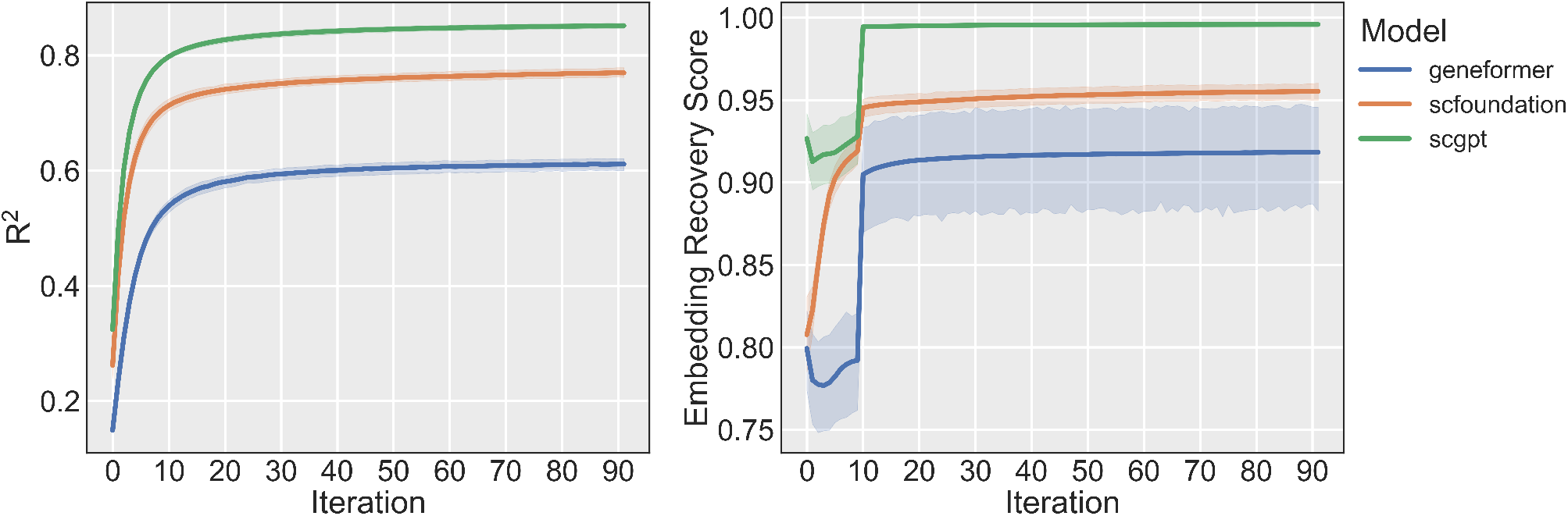
The training metrics *R*^2^ and Embedding Recovery Score for SAEs trained on the pre-trained models and the Covid-19 dataset. Lines represent the mean across the 12 layers, and shaded regions indicate standard deviation.

### A.6. Feature characterization

This appendix provides detailed examples of interpretable features discovered by sparse autoencoders across different models and datasets. Features are organized into two main categories as described in Section 3.2: gene-specific features that reflect properties of individual genes, and cell-specific features that capture properties of entire cells through contextual information learned by the transformer models.

#### A.6.1. Gene-specific features

##### Expression Level

These features activate within specific ranges of gene expression values. To quantify this relationship, we computed the Spearman rank correlation between feature activations and input gene expression vectors, after centering by the mean expression across all positions where the feature is active. Note that scGPT uses binned expression values (1-51), while scFoundation uses continuous values (0-8.77 for the COVID-19 dataset).

**Table A.2.** Expression level features in pre-trained scGPT (COVID-19 dataset)

| Feature | Expression (mean ± SD) | Spearman correlation | Density |
| --- | --- | --- | --- |
| 403 | 41.38 ± 4.76 | 0.58 | 0.18 |
| 227 | 32.16 ± 6.37 | 0.41 | 0.15 |
| 398 | 14.23 ± 6.55 | 0.44 | 0.16 |
| 92 | 5.48 ± 4.38 | 0.65 | 0.18 |

**Table A.3.**
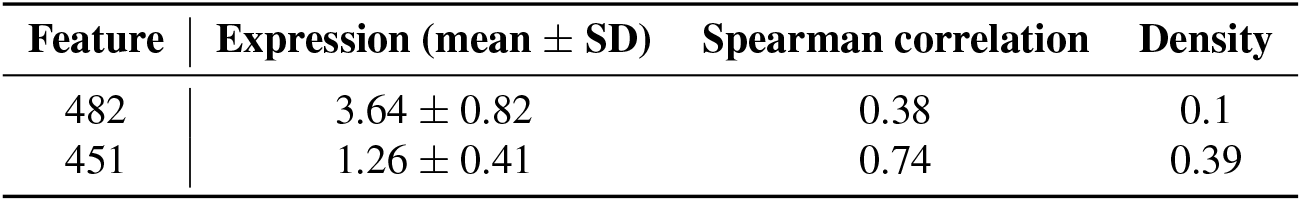
Expression level features in pre-trained scFoundation (COVID-19 dataset)

| Feature | Expression (mean ± SD) | Spearman correlation | Density |
| --- | --- | --- | --- |
| 482 | 3.64 ± 0.82 | 0.38 | 0.1 |
| 451 | 1.26 ± 0.41 | 0.74 | 0.39 |

##### Positional

Geneformer models the relative expression of genes with their position within the sequence. A SAE trained on Geneformer will learn positional features that have a stronger activation on different parts of the sequence.

**Figure A.4.**
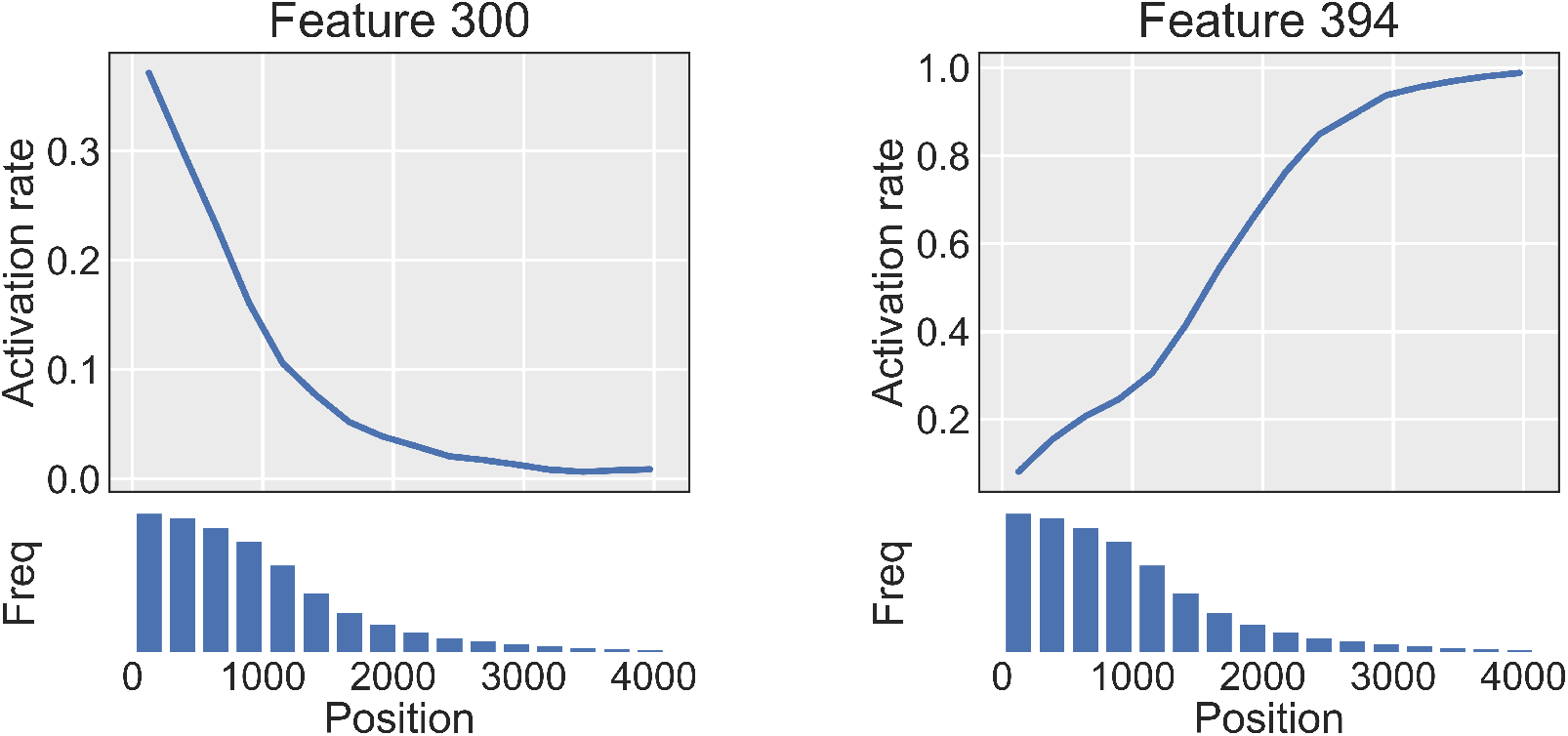
The activation rate of tokens at different positions in the gene sequence, normalized by the frequency a valid (non-padding) token appears at that position. Shown for features 300 and 394.

##### Gene family

These are features that activate specifically for genes belonging to particular functional families. Feature activations were thresholded at 0.3 to identify the most specific patterns.

**Table A.4.** Gene family features in pre-trained scGPT (COVID-19 dataset).

| Feature | Gene family | Percentage (%) | Unique Genes |
| --- | --- | --- | --- |
| 287 | Immunoglobulins | 92 | 57 |
| 334 | Metallothioneins | 100 | 4 |
| 276 | Histones | 95 | 26 |
| 302 | T cell receptors | 40 | 41 |
| 262 | Human leukocyte antigens | 99 | 13 |
| 172 | Mitochondrially encoded protein genes | 100 | 25 |
| 181 | Ribosomal protein genes | 95 | 79 |

**Table A.5.**
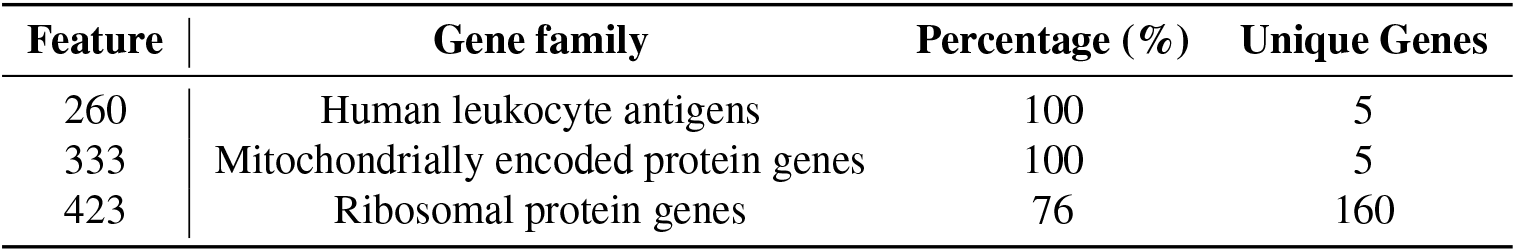
Gene family features in pre-trained scFoundation (COVID-19 dataset).

| Feature | Gene family | Percentage (%) | Unique Genes |
| --- | --- | --- | --- |
| 260 | Human leukocyte antigens | 100 | 5 |
| 333 | Mitochondrially encoded protein genes | 100 | 5 |
| 423 | Ribosomal protein genes | 76 | 160 |

**Table A.6.** Gene family features in fine-tuned Geneformer (COVID-19 dataset).

| Feature | Gene family | Percentage (%) | Unique Genes |
| --- | --- | --- | --- |
| 483 | Human leukocyte antigens | 81 | 14 |
| 125 | Mitochondrially encoded protein genes | 100 | 10 |
| 38 | Histones | 82 | 67 |

##### Biological process

These features capture genes involved in specific biological processes through functional gene modules. Feature activations were thresholded at above 0.5 to identify the strongest patterns, and enrichment is reported as the adjusted p-value.

**Table A.7.** Biological process features in pre-trained scGPT (COVID-19 dataset).

| Feature | Biological process | Adj. p-value |
| --- | --- | --- |
| 50 | Nuclear Outer Membrane-ER Network | 9.00e-48 |
| 173 | Cell Cycle | 1.70e-92 |
| 213 | Phagocytosis | 8.43e-18 |
| 233 | Hemostasis | 1.52e-21 |
| 330 | Adaptive Immune Response | 5.56e-60 |

**Table A.8.** Biological process features in pre-trained scGPT. Here the SAE was trained on the CellxGene Census and evaluated across CellxGene and COVID-19 datasets.

| Feature | Biological process | Adj. p-value for COVID-19 | Adj. p-value for CellxGene |
| --- | --- | --- | --- |
| 114 | Cell Cycle | 2.58e-88 | 1.86e-210 |
| 404 | ER to Golgi Transport | 2.101e-34 | 1.12e-73 |
| 765 | Programmed Cell Death | 3.27e-12 | 3.73e-79 |
| 816 | Response to Virus | 1.88e-27 | 1.75e-96 |

**Table A.9.** Biological process features in finetuned Geneformer (COVID-19 dataset).

| Feature | Biological process | Adj. p-value |
| --- | --- | --- |
| 279 | Cell Cycle | 5.27e-131 |
| 75 | Nuclear Outer Membrane-ER Network | 1.43e-40 |
| 201 | Programmed Cell Death | 2.59e-19 |
| 86 | Adaptive Immune Response | 7.94e-37 |
| 429 | Hemostasis | 4.10e-39 |

#### A.6.2. Cell specific

##### Cell type

These features correspond to specific cell types, evaluated using Adjusted Mutual Information (AMI) and F1 scores with optimized activation thresholds per cell type.

**Table A.10.**
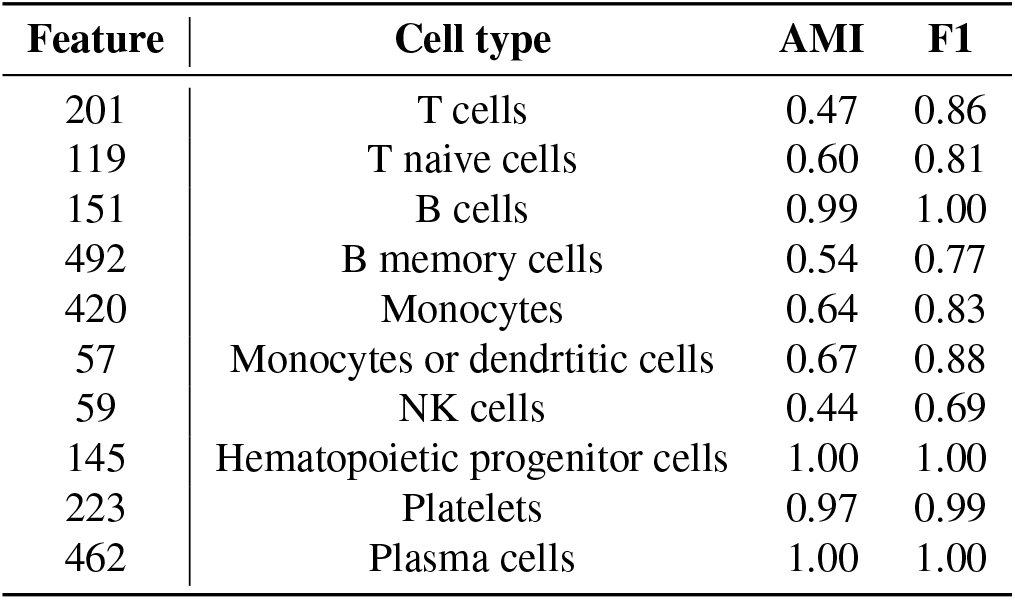
Cell type features in pre-trained scGPT (COVID-19 dataset)

**Table A.11.** Cell type features in pre-trained scFoundation (COVID-19 dataset)

| Feature | Cell type | AMI | F1 |
| --- | --- | --- | --- |
| 266 | T cells | 0.32 | 0.79 |
| 194 | T naive cells | 0.56 | 0.79 |
| 190 | B cells | 0.93 | 0.98 |
| 445 | B memory cells | 0.50 | 0.75 |
| 277 | Monocytes | 0.58 | 0.80 |
| 230 | Monocytes or dendritic cells | 0.61 | 0.83 |
| 380 | NK cells | 0.34 | 0.62 |
| 508 | Hematopoietic progenitor cells | 0.30 | 0.50 |
| 116 | Platelets | 0.88 | 0.95 |
| 337 | Plasma cells | 0.51 | 0.69 |

**Table A.12.** Cell type features in fine-tuned Geneformer (COVID-19 dataset)

| Feature | Cell type | AMI | F1 |
| --- | --- | --- | --- |
| 454 | T cells | 0.45 | 0.85 |
| 419 | T naive cells | 0.40 | 0.62 |
| 233 | B cells | 0.99 | 1.00 |
| 19 | B memory cells | 0.45 | 0.65 |
| 173 | Monocytes | 0.79 | 0.91 |
| 121 | Monocytes or dendritic cells | 0.86 | 0.98 |
| 156 | NK cells | 0.65 | 0.85 |
| 456 | Hematopoietic progenitor cells | 0.35 | 0.5 |
| 325 | Platelets | 1.00 | 1.00 |
| 77 | Plasma cells | 1.00 | 1.00 |

##### Disease

Features that preferentially activate in cells from patients with specific disease conditions, evaluated using Adjusted Mutual Information (AMI) and F1 scores with optimized activation thresholds.

**Table A.13.** COVID-19 related features in pre-trained scGPT (COVID-19 dataset)

| Feature | Disease | AMI | F1 |
| --- | --- | --- | --- |
| 127 | COVID-19 | 0.11 | 0.59 |
| 89 | post-COVID-19 disorder | 0.20 | 0.55 |

**Table A.14.** COVID-19 related features in pre-trained scFoundation (COVID-19 dataset)

| Feature | Disease | AMI | F1 |
| --- | --- | --- | --- |
| 186 | COVID-19 | 0.21 | 0.63 |
| 250 | post-COVID-19 disorder | 0.18 | 0.50 |

##### Sequencing Technology

Features that activate based on technical aspects of the data, including sequencing technologies and experimental protocols, evaluated using Adjusted Mutual Information (AMI) and F1 scores with optimized activation thresholds.

**Table A.15.**
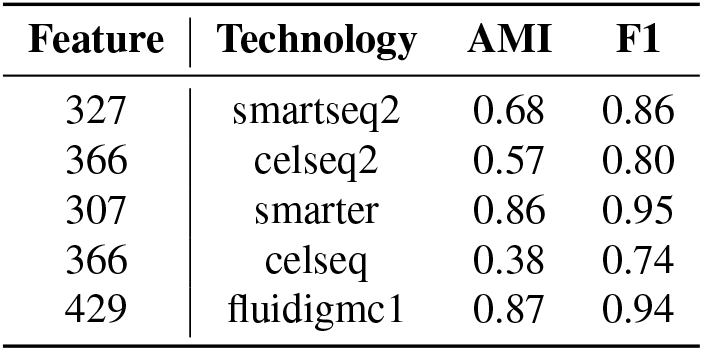
Sequencing technology features in pre-trained scGPT (Pancreas dataset)

**Table A.16.**
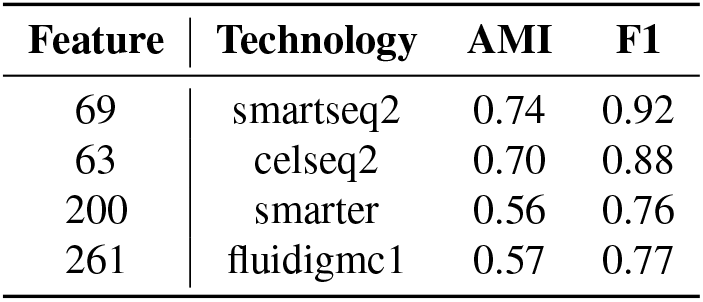
Sequencing technology features in pre-trained scFoundation (Pancreas dataset)

#### A.6.3. Specific feature examples

**Figure A.5.**
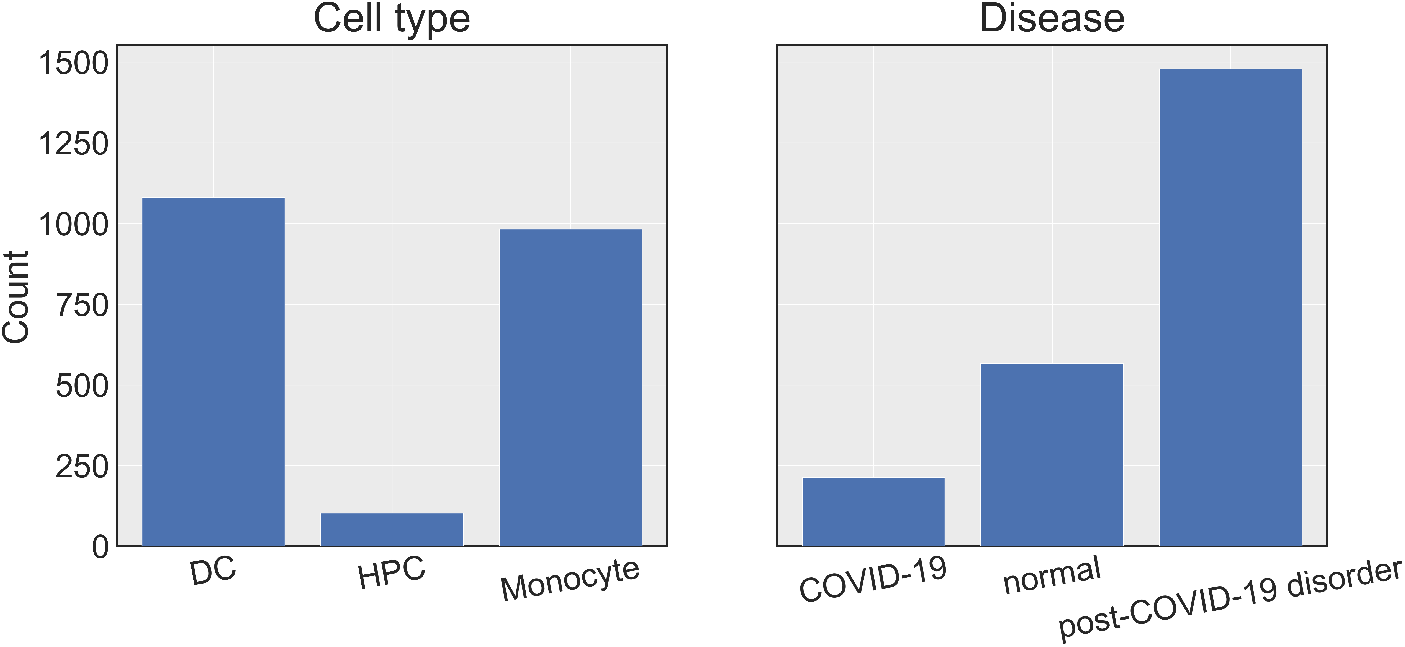
Distribution of token activations of feature 224 across cell types and COVID-19 status. Feature activations were thresholded above 0.5

##### A.6.4. CellXGene batch effects

**Table A.17.** Batch features for pre-trained scGPT on CellXGene Census data. Batch effects include technical variations from sequencing technologies and original studies (DS1–DS6 refer to datasets detailed in Table S18).

| Feature | Batch | AMI | F1 |
| --- | --- | --- | --- |
| 179 | DS1 | 0.63 | 0.74 |
| 337 | DS2 | 0.68 | 0.76 |
| 338 | DS3 | 0.68 | 0.76 |
| 341 | DS4 | 0.76 | 0.86 |
| 540 | DS5 | 0.92 | 0.96 |
| 832 | DS6 | 0.89 | 0.94 |
| 701 | Seq-Well | 0.31 | 0.44 |

**Table A.18.**
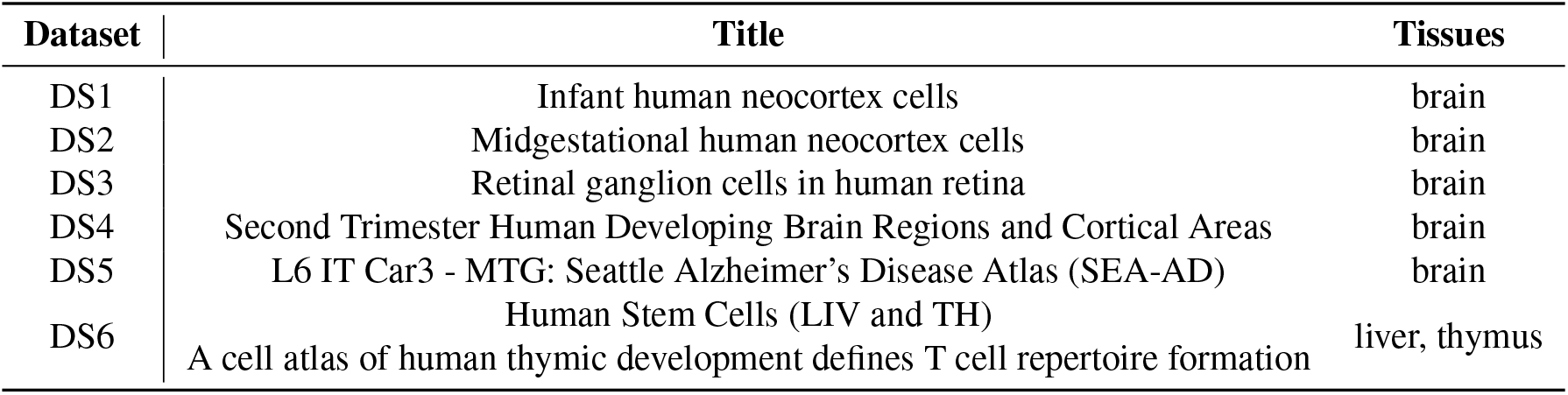
Characteristics of some datasets from the CellXGene Census. Note that brain tissues are overrepresented due to the composition of the CellXGene Census itself, where about half of the available datasets focus on brain-related samples.

**Figure A.6.**
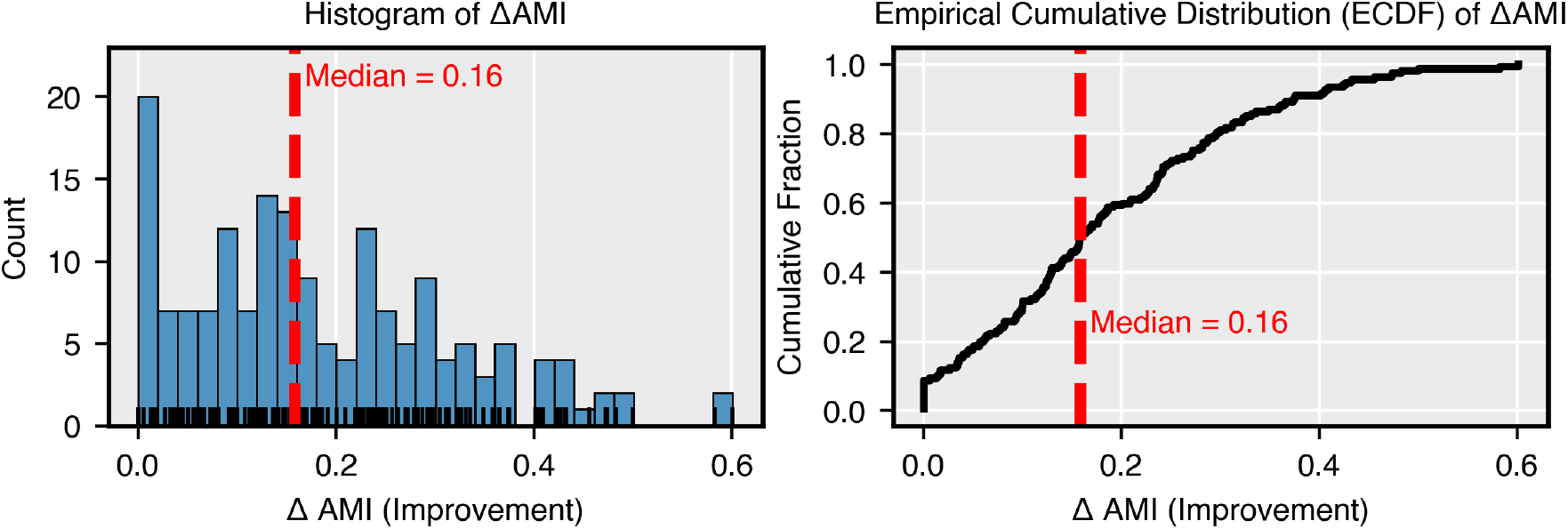
Distribution of the ΔAMI calculated between feature activations and two cell type annotations. Values above zero indicate features that better capture cell types within specific studies than across all studies.

##### A.6.5. Cell type features over model layers

**Figure A.7.**
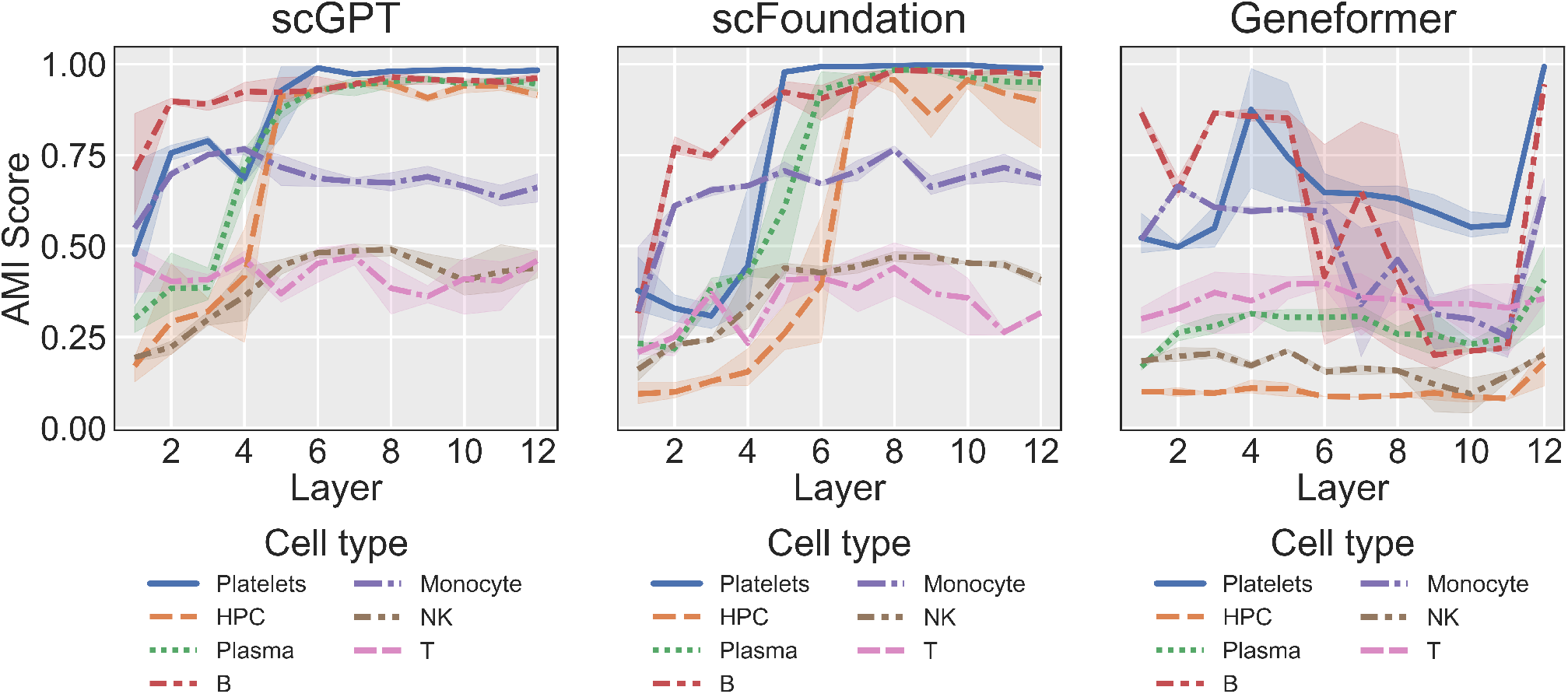
Layer-wise analysis of cell type feature emergence in pre-trained scGPT, scFoundation and Geneformer. For each layer, we extracted the feature most associated with each cell type by their AMI score. Each line represents a different cell type of the corresponding model. This analysis was conducted on the Covid-19 dataset.

### A.7. Batch steering

**Figure A.8.**
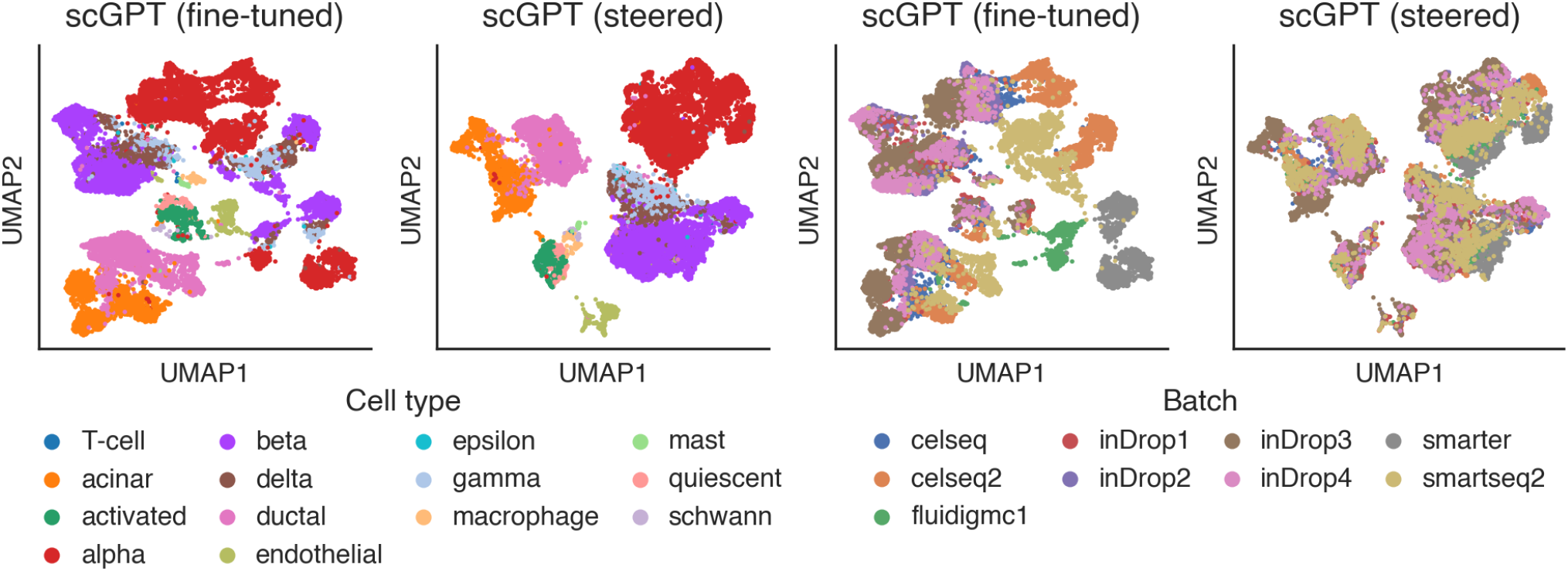
UMAP of the standard and steered cell embeddings of fine-tuned scGPT, colored by cell type (left) and sequencing protocol (right).

**Figure A.9.**
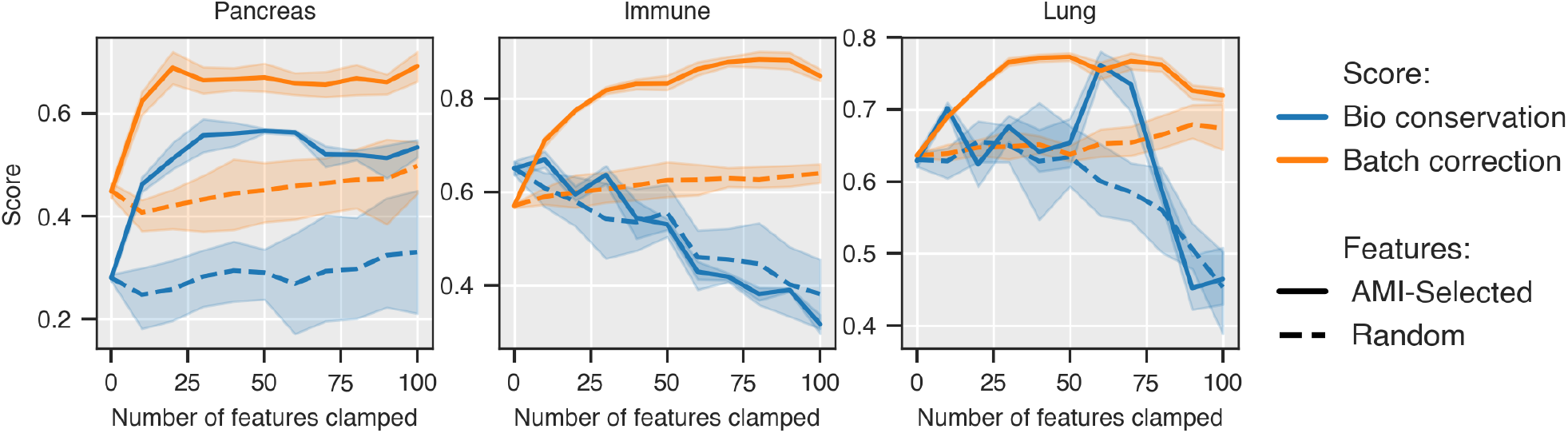
Biological conservation and batch correction scores as features are sequentially steered, selected randomly or by maximum AMI, across three datasets. Lines show the mean over five seeds, and shaded regions indicate standard deviation. The sparse autoencoders were trained on the fine-tuned scGPT model.

**Figure A.10.**
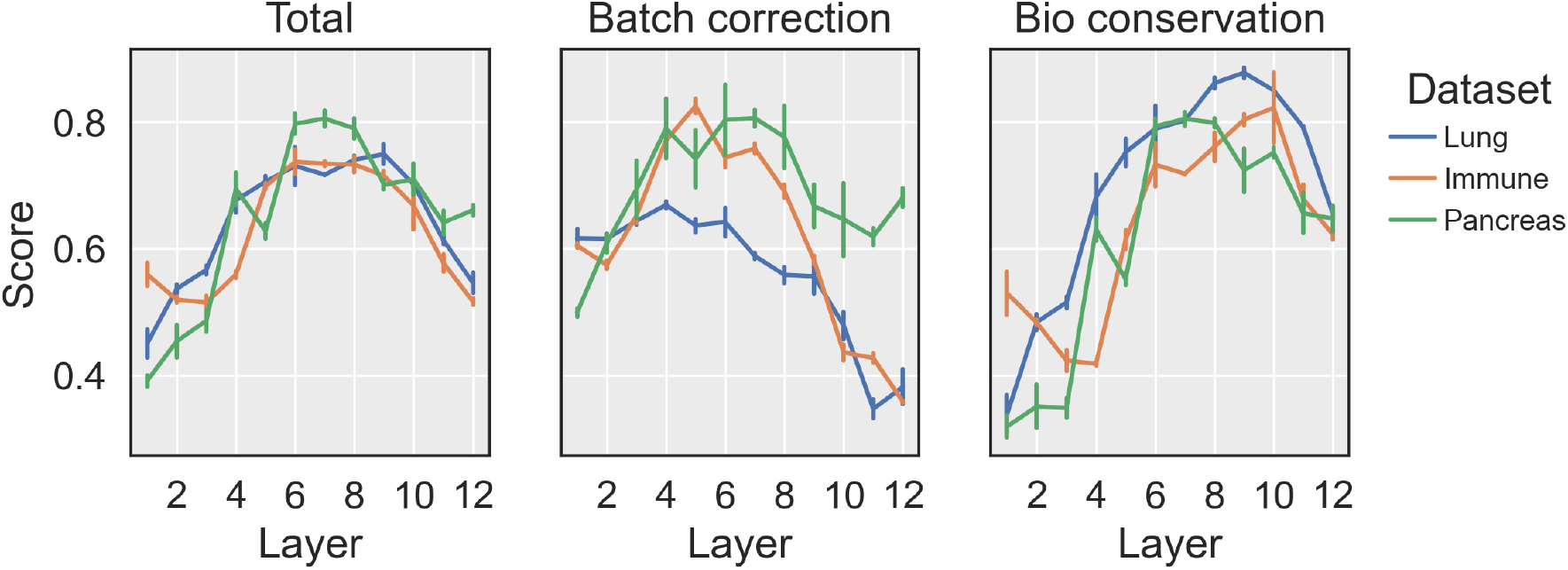
Layer-wise performance of SAE feature steering for batch correction. Performance metrics (total score, batch correction, and biological conservation) are shown for features extracted from different layers (1-12) of scGPT across three datasets: Lung (blue), Immune (orange), and Pancreas (green). Error bars represent standard deviation across five random seeds.

**Figure A.11.**
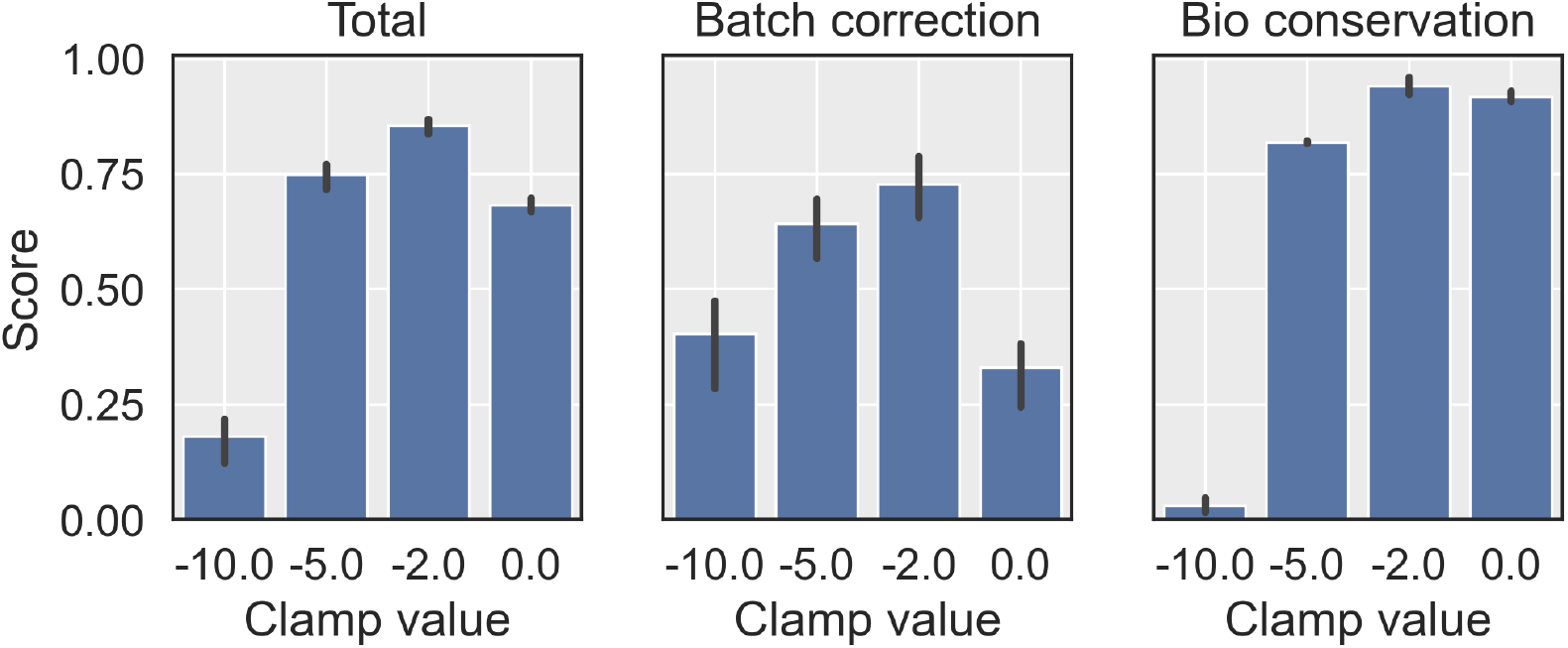
Effect of different clamping values on batch correction performance for scGPT. Comparison of different clamping values (−10.0, −5.0, −2.0, 0.0) on three evaluation metrics: total score (left), batch correction score (middle), and biological conservation score (right). Bar heights represent mean scores across datasets, with error bars indicating standard deviation.

**Figure A.12.**
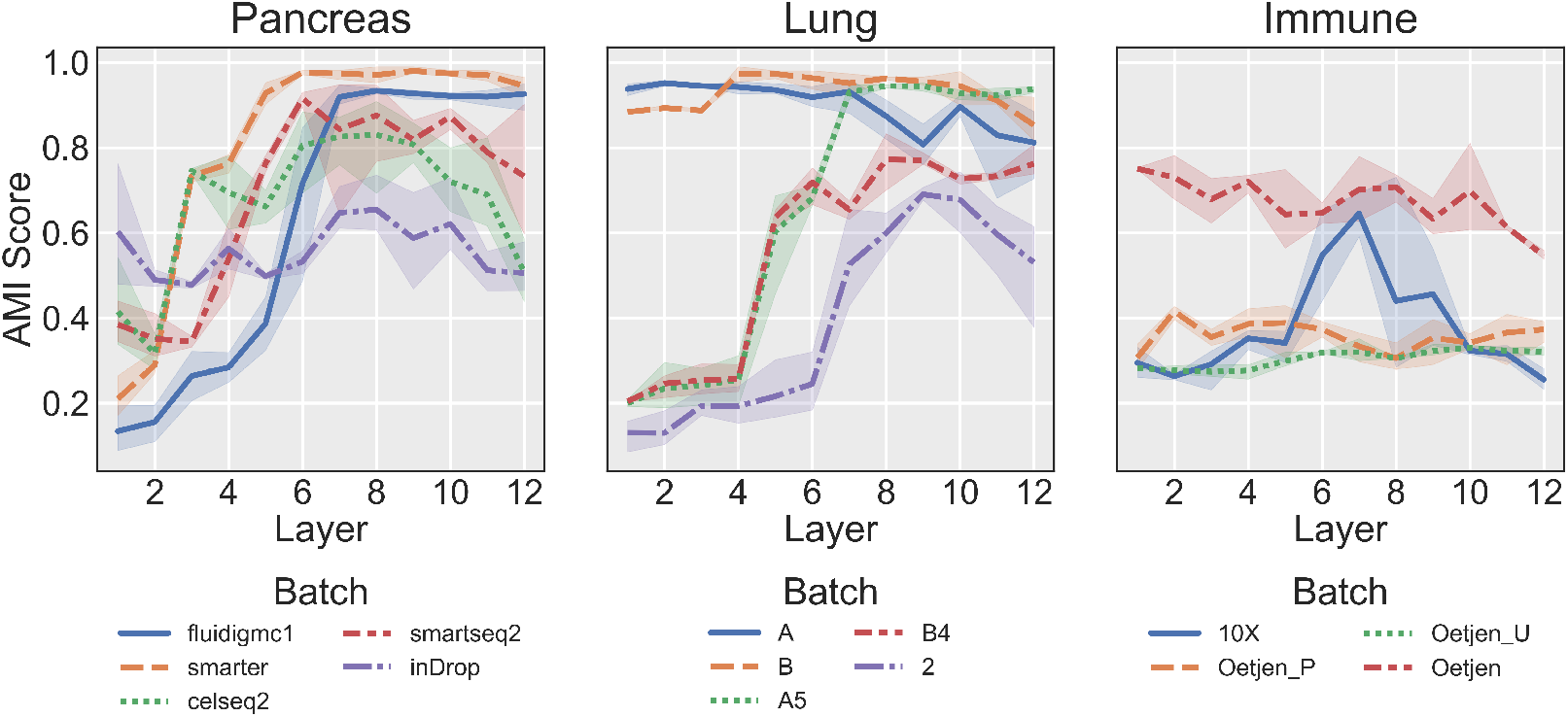
Layer-wise analysis of batch feature emergence in fine-tuned scGPT. For each layer, we extracted the feature most associated with each batch effect by their AMI score. Each line represents a different batch variable of the corresponding dataset.

**Table A.19.** Batch correction performance of single-cell foundation models on the lung dataset. Values show batch correction, biological conservation, and total scores, with higher scores indicating better performance. Means and standard deviations are computed across five runs with different random model initializations. Bold values highlight the best method within each category.

| Method | Batch correction | Bio conservation | Total |
| --- | --- | --- | --- |
| Unintegrated (PCA) | 0.22±0.000 | 0.65±0.014 | 0.48±0.008 |
| scVI | 0.86±0.010 | 0.54±0.056 | **0.67±0.003** |
| scFoundation (zero-shot) | 0.51±0.000 | 0.59±0.002 | 0.56±0.001 |
| scFoundation (zero-shot, steered) | 0.53±0.000 | 0.60±0.008 | **0.57±0.005** |
| Geneformer (fine-tuned) | 0.49±0.000 | 0.99±0.001 | 0.79±0.001 |
| Geneformer (fine-tuned, steered) | 0.61±0.00 | 0.98±0.003 | **0.83±0.002** |
| scGPT (zero-shot) | 0.65±0.009 | 0.08±0.003 | **0.31±0.004** |
| scGPT (zero-shot, steered) | 0.59±0.025 | 0.01±0.002 | 0.24±0.011 |
| scGPT (fine-tuned) | 0.56±0.006 | 0.62±0.016 | 0.59±0.011 |
| scGPT (fine-tuned, DAR) | 0.68±0.003 | 0.58±0.005 | 0.62±0.003 |
| scGPT (fine-tuned, steered) | 0.71±0.003 | 0.62±0.009 | **0.65±0.004** |

**Table A.20.** Batch correction performance of single-cell foundation models on the immune dataset. Values show batch correction, biological conservation, and total scores, with higher scores indicating better performance. Means and standard deviations are computed across five runs with different random model initializations. Bold values highlight the best method within each category.

| Method | Batch correction | Bio conservation | Total |
| --- | --- | --- | --- |
| Unintegrated (PCA) | 0.11±0.000 | 0.55±0.004 | 0.37±0.002 |
| scVI | 0.74±0.008 | 0.73±0.016 | **0.73**±0.011 |
| scFoundation (zero-shot) | 0.47±0.006 | 0.57±0.04 | **0.53**±0.026 |
| scFoundation (zero-shot, steered) | 0.44±0.02 | 0.44±0.084 | 0.44±0.040 |
| Geneformer (fine-tuned) | 0.52±0.000 | 0.91±0.002 | 0.76±0.001 |
| Geneformer (fine-tuned, steered) | 0.60±0.00 | 0.92±0.017 | **0.79**±0.01 |
| scGPT (zero-shot) | 0.66±0.011 | 0.11±0.005 | **0.33**±0.005 |
| scGPT (zero-shot, steered) | 0.78±0.018 | 0.02±0.015 | 0.32±0.012 |
| scGPT (fine-tuned) | 0.52±0.004 | 0.61±0.010 | 0.58±0.005 |
| scGPT (fine-tuned, DAR) | 0.75±0.006 | 0.57±0.017 | 0.64±0.011 |
| scGPT (fine-tuned, steered) | 0.72±0.010 | 0.60±0.012 | **0.65**±0.005 |

### A.8. Drug concentration steering

**Figure A.13.**
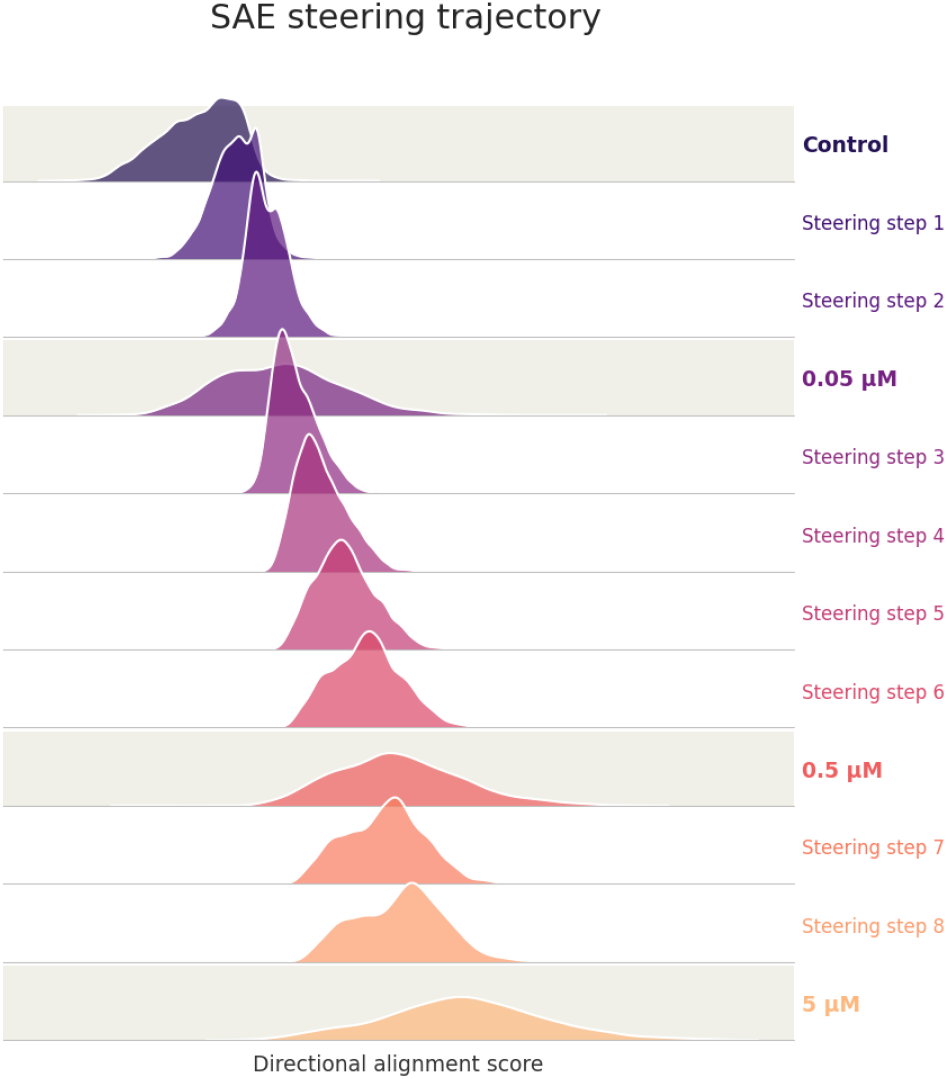
Distributions of the centroid projection score at each steering step, alongside the real distributions for cells treated at 0.05 µM, 0.5 µM, and 5 µM Panobinostat (highlighted rows).

## Footnotes

1 https://anonymous.4open.science/r/sae-for-scFMs-ED25/.

